# Taxonomy, distribution and host relationships of aphidiine wasps (Hymenoptera: Aphidiidae) parasitizing aphids (Hemiptera: Aphididae) in Australian grain production landscapes

**DOI:** 10.1101/2021.01.12.426457

**Authors:** Samantha Ward, Paul A. Umina, Andrew Polaszek, Ary A. Hoffmann

## Abstract

Aphid parasitoids (Hymenoptera; Aphidiidae) were surveyed within grain production landscapes in Victoria, Australia between 2017 and 2018, as well as more sporadically nationwide between 2016 and 2019. In addition, aphidiine records were collated from insect depositories around Australia and online databases. The 5551 specimens recorded constituted a total of 23 species and seven genera. *Diaeretiella rapae* (M’Intosh) was the most common species, representing more than 70% of all aphidiines recorded. This species also showed a greater northerly geographic range than other aphidiines. During sampling between 2017 and 2019, aphidiines were reared from mummies to ascertain host-parasitoid relationships. *Diaeretiella rapae* was again the most commonly reared parasitoid, although aphidiine preference varied with aphid host and between states and territories. An illustrated dichotomous key to Australian aphidiines in grain production landscapes is provided for the 11 species sampled in our field surveys. This is the first comprehensive review of aphidiines sampled within Australia in over two decades. Knowledge about the diversity and distribution of these parasitoids is important for understanding their impact on current and future invasions of aphid species. In addition, understanding the interactions between grain aphids and their associated parasitoids will further support the inclusion of parasitoid wasps into integrated pest management (IPM) strategies.

## 1. Introduction

Parasitoid Hymenoptera are a particularly abundant and species-rich group of organisms, numbering in the hundreds of thousands of species. The majority are directly beneficial in economic terms, frequently functioning as determinants of host population densities (Doutt 1959; Lasalle & Gauld 1991). As such, parasitoid Hymenoptera can be utilised in biological pest control programs. One of the first successful and best-known examples of this is the control of the greenhouse whitefly (*Trialeurodes vaporariorum* (Westwood)) with the parasitoid, *Encarsia formosa* Gahan. Commercially available in England by 1930, it was later exported to Canada, Australia, New Zealand and mainland Europe, where it was successful in the control of *T. vaporariorum* (Gerling 1966; Van Lenteren 1983). Over 216 species of invertebrate pests have been controlled, either in part or completely, by the introduction of parasitoids (Greathead 1986; Laing & Hamai 1976). Groups of parasitoid Hymenoptera most successful as biological control agents include the Aphelinidae and Braconidae (Greathead 1986), both relevant to aphid control.

Until the early 1990s, the aphidiines were treated as a separate family, the Aphidiidae, based on behavioural and morphological traits, until phylogenetic studies showed them to be a lineage within the Braconidae (Quicke & Van Achterberg 1990; Van Achterberg & Quicke 1992; Wharton et al. 1992). Aphidiinae are solitary obligate endoparasitoids of adult and immature aphids (Hemiptera: Aphididae), which are found worldwide (Starý 1970). The aphidiine larva progressively consumes the soft tissue of the aphid host as it develops inside it, and then spins a cocoon within the host integument (or underneath in the case of the genus *Praon*), forming a characteristic ‘mummy’ (Belshaw & Quicke 1997).

Several species of aphidiine wasps have been successfully used in biological control programs around the world (Carver 1989). Within Australia, however, little is known about the diversity and distribution of Aphidiinae, particularly in the context of parasitizing grain aphids. Carver and Starý (1974) surmised, based on their knowledge of the aphid fauna of Australia, that the aphidiid fauna would be small and most likely introduced, with the number of species fewer than expected based on surveys in other countries. These authors noted the presence of mostly introduced aphidiines, including *Aphidius platensis* Brèthes, *Aphidius salicis* Haliday, *Diaeretiella rapae* (M’Intosh), *Ephedrus persicae* Froggatt, *Paraphedrus relictus* Stary and Carver, *Trioxys cirsii* (Curtis), *Trioxys pallidus* (Haliday), and *Trioxys tenuicaudus* Stary, in addition to some other unidentified species within Australia. However, Carver and Starý (1974) noted difficulties identifying species of *Trioxys*, with the closely related species identified so far likely an underestimate of the current diversity within Australia. More recently, Heddle et al. (2020) monitored aphidiines associated with the recently arrived Russian wheat aphid (*Diuraphis noxia* (Mordvilko ex Kurdjumov)) over one growing season and discovered eight aphidiine species parasitizing this aphid pest in Australia.

Constant co-operation is required between taxonomists and biological control specialists for successful biological control programs (de Moraes 1987). More specifically, the success of using aphidiines as biological control agents is reliant upon understanding their taxonomy, ecology, and host selection behaviour (see Rehman and Powell (2010)). Understanding ecological interactions and geographical species distributions is fundamental to recognising biodiversity patterns in parasitoids, as well as the abiotic and biotic processes that affect parasitoid populations (Rushton et al. 2004). In addition, mapping species distributions at a spatial scale is required for regional biodiversity conservation planning (Ferrier et al. 2002). Distribution mapping of aphid parasitoids is particularly useful from a biological control viewpoint because it helps to identify species that have the ability to tolerate wide ranges of climates and habitats in helping to deduce whether they have the capacity to establish within a particular region (Gonzalez et al. 1978).

The Aphidiinae is a subfamily of parasitoid wasp found within the family Braconidae (Insecta: Hymenoptera). The subfamily is diverse, containing 50 genera and approximately 400 species (Mackauer & Starý 1967; Starý 1988), although this latter figure is likely an underestimate. Despite in-depth assessments of their morphology, the phylogenetic relationships between Aphidiinae and other braconid subfamilies are poorly understood (Shaw & Huddleston 1991; Shi & Chen 2005; Wharton 1993). The most widely accepted classification of the Aphidiinae involves their subdivision into four tribes (Aphidiini, Aclitini, Praini, and Ephedrini), as described by Mackauer (1961). Belshaw and Quicke (1997) explain this distinction in greater detail. The tribal classification has been disputed by authors such as Sanchis et al. (2000) who sequenced 37 aphidiine taxa using 18S rDNA. Although two of the four traditionally accepted tribes (Ephedrini and Praini) were confirmed, two others were questioned. Later analysis suggested instead that there might be three tribes (Ephedrini, Praini, and Aphidiini) or that a new classification system of five or more tribes (Ephedrini, Praini, Monoctonini, Trioxini, and Aphidiini) is required (Sanchis et al. 2000).

Aphidiinae keys are currently available from around the world, however none of these focus on Australian fauna. A few examples of studies contributing relevant information include those undertaken in Iran (Rakhshani et al. 2012; Rakhshani et al. 2008b), North America (Pike et al. 1997), south-eastern Europe (Kavallieratos et al. 2010), and Malta (Rakhshani et al. 2015). Early descriptions of endemic Australian ichneumonids were provided by J. C. Fabricius in 1775 from specimens collected by Sir Joseph Banks, who accompanied Captain James Cook on his voyage around Australia (1768-1771) (Gauld 1984), but limited work has been done since that time.

The aim of this paper was to classify aphidiine wasps reared from grain aphids within Australia through a combination of morphological identification and CO1 barcoding. To understand the geographic ranges of each species, distribution maps were created. Furthermore, tri-trophic relationships were considered between host plant, host aphid, and aphidiine to ascertain the host range of each species within grains production landscapes.

## 2. Methodology

### 2.1. Field collections

#### 2.1.1. Repeat sampling in Victoria, Australia (2017-2018)

Repeat sampling occurred in Victoria (VIC), Australia, between 2017 and 2018. Crop types included canola, wheat, barley, and a single field containing a cover crop of wheat and forage brassicas (Fig. 1).

**Figure 1:**
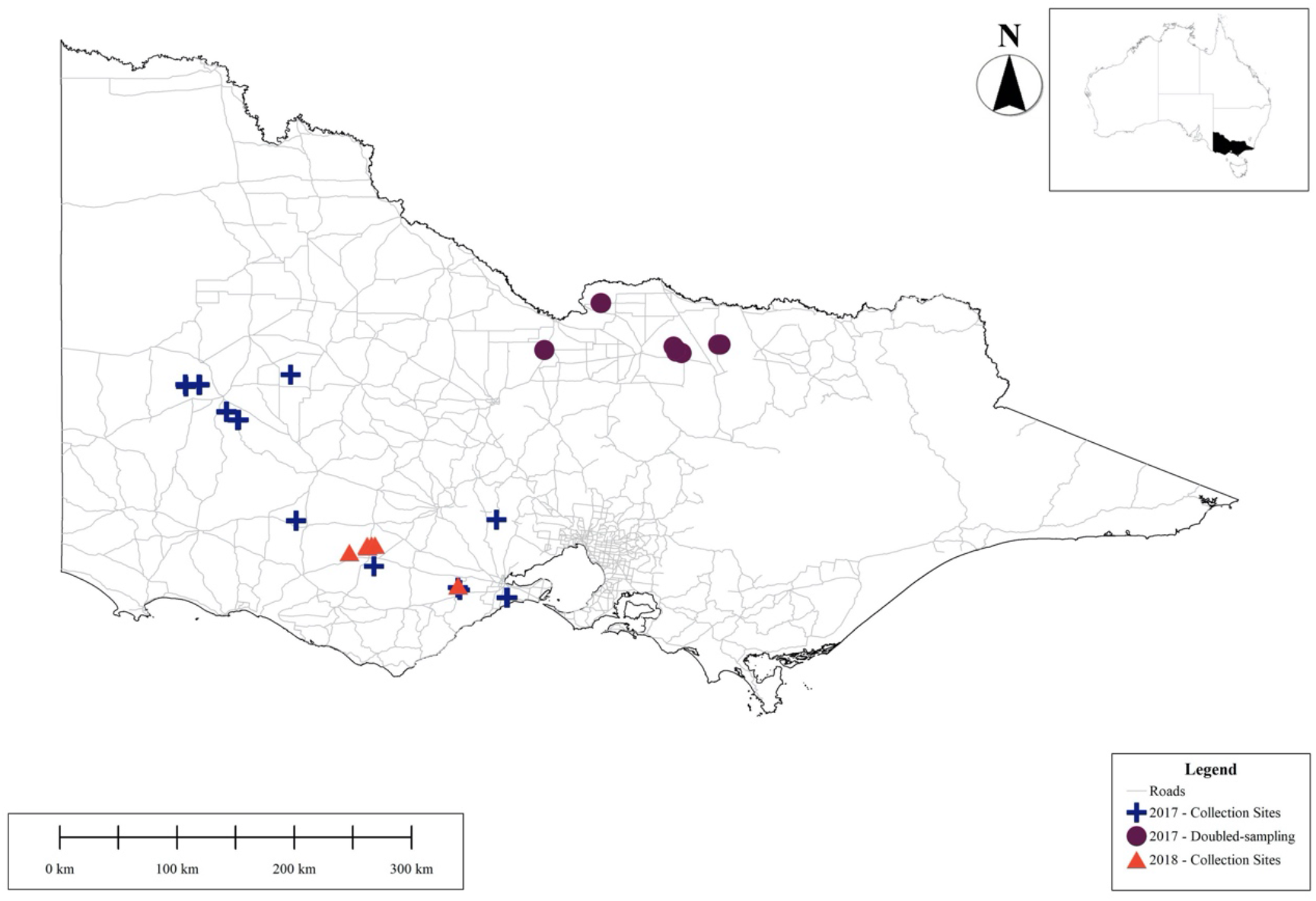
Map of 2017 and 2018 repeat sampling sites in Victoria, Australia (main map depicts sites at state-level, inset map depicts the state of Victoria) (Department of Agriculture and Water Department of Agriculture and Water Resources 2020; Public Service Mapping Agency 2020).

A total of 84 sampling trips were undertaken in 2017, with 16 fields sampled every 6-12 weeks (either three or six times during the season, with a further six sampled twice throughout the season. Sites sampled six times are referred to as ‘doubled-sampling’ sites in Fig. 1. Parasitoids were sampled using yellow pan traps and through direct searching and rearing from mummies in 2017, both within the crops and neighbouring vegetation. In 2018, six fields (three canola and three wheat) were sampled 10 times each. Parasitoids were directly sampled and reared from mummies from each field and neighbouring vegetation every three weeks. Additionally, parasitoids were vacuum sampled every six weeks within each field and neighbouring vegetation. Further details of sampling are provided in Ward et al. (2021).

#### 2.1.2. Repeat sampling in Australia (2019)

Repeat sampling was undertaken within 26 canola paddocks every four weeks throughout the growing season in 2019, across four Australian states: New South Wales (NSW), South Australia (SA), VIC, and Western Australia (WA) (Fig. 2).

**Figure 2:**
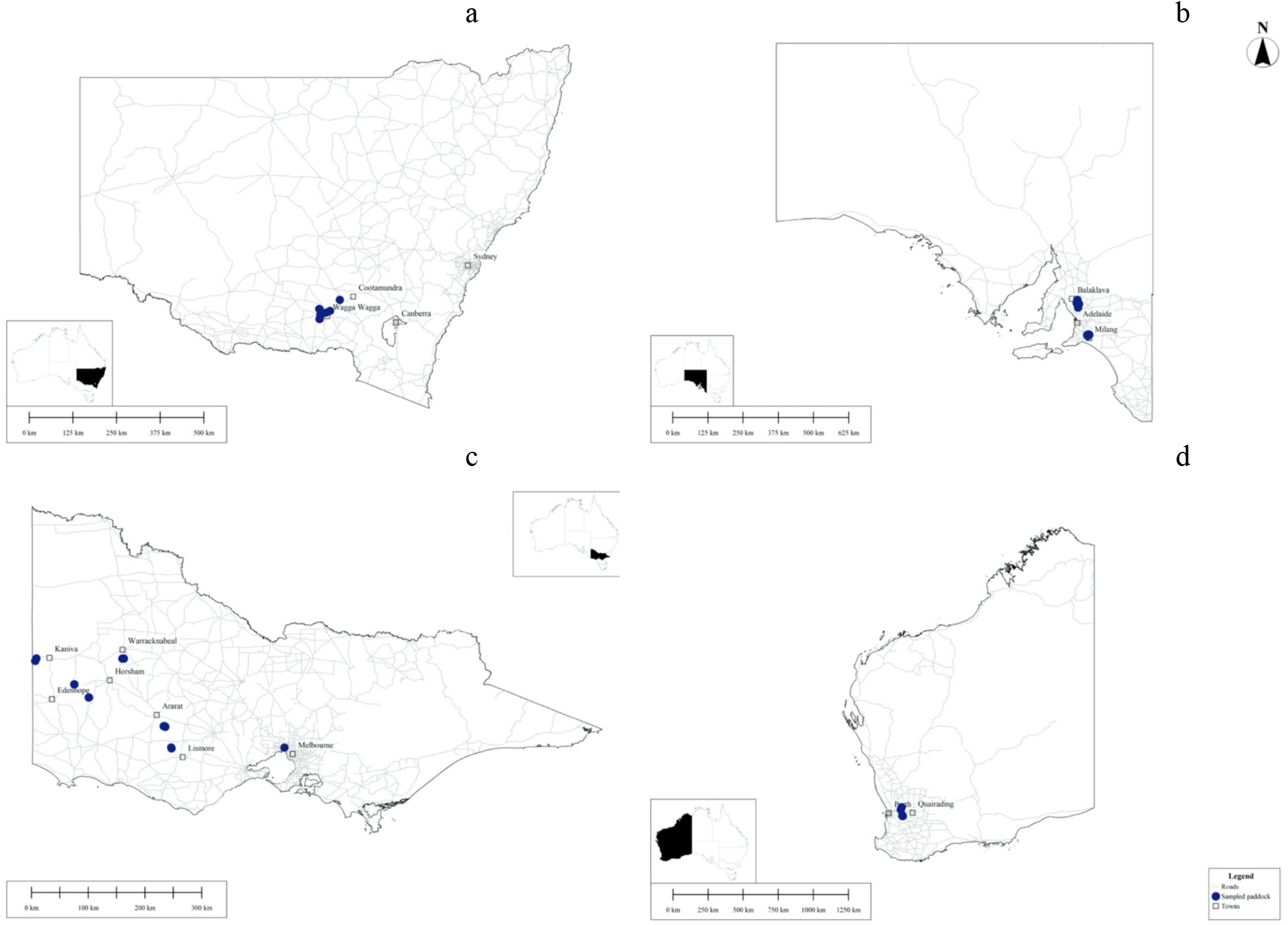
Map of 2019 repeat sampling sites in Australia for each state: a) NSW, b) SA, c) VIC, d) WA (inset maps depict states on Australian map) (Department of Agriculture and Water Department of Agriculture and Water Resources 2020; Public Service Mapping Agency 2020).

Parasitoids were reared from directly sampled mummies. Sampling began at the edge of the field and continued into the paddock, with sampling points spaced at least 30 m apart. *Myzus persicae* (Sulzer) was targeted, where possible. Further details are provided in Ward et al. (2021).

#### 2.1.3. Opportunistic sampling in Australia (2016-2019)

Between 2016 and 2019, opportunistic collections of mummified aphids and their associated hymenopteran primary and secondary parasitoids were made in grain crops on a national scale by volunteers, using a variety of trapping methods. Methods included direct searching, sweep netting, vacuum sampling, pan trapping and rearing. The information collected alongside the samples included the trapping method, GPS co-ordinates and location data, collection date, crop type, host plant, and collector. Grain crops were prioritised, however occasionally horticultural crops were sampled to draw comparisons between species composition. *Myzus persicae* was targeted during the 2019 sampling period. Parasitoid specimens were received at the laboratory in 70-100% ethanol for morphological identification. In addition, mummies on plant material were received and parasitoids were reared from them.

Mummies were transferred onto petri dishes made up with a 1% agar solution, within which 2-3 cotyledons of sprouting radish (*Raphanus raphanistrum* subsp. *sativus* L.), cuttings of wheat (*Triticum aestivum* L.), or the plant material on which the mummies were collected, were inserted. The plant material selected pertained to the host plant from which the mummies were collected. Mummies from oilseeds were reared on *R. raphanistrum* and mummies from cereals were reared on *T. aestivum*. Petri dishes were sealed with a lid lined with filter paper. Mummies were placed onto the plant material and each dish was placed in a constant temperature cabinet maintained at 20°C with a 16L:8D photoperiod. Leaves were changed weekly, or immediately once the plants began to look limp or discoloured, if fungus growth was evident, or if the filter paper became moist.

Due to the paucity of samples from Tasmania, a trip was undertaken to this state in October 2019, surveying roadside verges and grain crops for the presence of aphids and associated parasitoids. We used both direct sampling of aphidiines and direct sampling of mummies to subsequently rear parasitoids. The rearing protocol followed that outlined above.

### 2.2. Specimens from insect depositories

Aphid parasitoids were provided by the individuals and insect collections listed in Table 1. Specimen data was recorded based on attached labels and, where specimens had incomplete taxonomic data, identifications were undertaken based on morphological analysis where possible. These specimens provided an overview of historic aphidiine collections within Australia. Where there was no provision of GPS information, location was calculated based on searching the name of the location and its description with reference to Google Maps.

**Table 1:** Individuals and insect depositories from which aphidiines were sourced (acronyms pertain to the relevant insect depository)

| **Insect depository acronym** | **Insect depository name, address** | **Individual providing specimens** |
| --- | --- | --- |
| ANIC | Australian National Insect Collection, CSIRO, Canberra, Australian Capital Territory | J. Rodriguez |
| ASCU | Agricultural Scientific Collections Unit, Orange Agricultural Institute, Orange, NSW | P. Gillespie |
| NMVM | National Museum of Victoria, Melbourne, VIC | K. Walker |
| QMBA | Queensland Museum, Brisbane, QLD | C. Burwell |
| UQIC | University of Queensland Insect Collection, St Lucia, QLD | C. Burwell |
| VAIC | Victorian Agricultural Insect Collection, Department of Primary Industries, Knoxfield, VIC | L. Semeraro |
| WARI | Waite Agricultural Research Institute, University of Adelaide, Glen Osmond, SA | A. Austin |

### 2.3. Morphological taxonomic classification & photography

All aphid parasitoids were sexed and morphologically identified with a Leica MS5 or Nikon SMZ1500 microscope, using keys produced by Rakhshani et al. (2015) and Rakhshani et al. (2012). Parasitoids were selected for further genetic identification when their morphology varied from any identifying features labelled within the keys or if specimens were in poor condition. Additionally, due to the variation in antennal segment numbers between sexes (Smith 1944), several males were barcoded, along with known females of particular species, to corroborate taxonomy. Sequences of a 658 base pair fragment of the cytochrome c oxidase subunit 1 gene (CO1) were compared for known species, with those of uncertain classification, such as cryptic taxa. Line drawings were made in Adobe Photoshop version 21.20, traced from photographs captured using a digital camera attached to a Nikon SMZ1500 microscope. SEM photography of specimens was undertaken using negatively stained grids observed on a FEI Tecnai F30 at the Bio21 Advanced Microscopy Facility and subsequently assembled in Adobe Photoshop version 21.20.

### 2.4. CO1 barcoding & phylogenetics

DNA was extracted using a modified non-destructive Chelex® extraction method, adapted from Walsh et al. (1991), as detailed in Carew et al. (2003). An individual wasp was placed within a 0.5ml micro-centrifuge tube, along with 3 μl of Proteinase K (20 mg/ml) and 70 μl of 5 % Chelex® solution, before being incubated in a 56°C water bath for 60 minutes. Afterwards, the tube was transferred to a 90°C water bath for 10 minutes, to ensure the sample breakdown was complete. Samples were homogenised in a TissueLyser (Qiagen, Hilden, Germany) for 2 min at 25 Hz, and after centrifuging, were incubated in a 65°C water bath for 60 minutes, before transferral to a 90°C water bath for 10 minutes. Prior to PCR, tubes were spun at 13,000-14,000 rpm for five minutes in a D3024 high-speed microcentrifuge (DLAB Scientific, Beijing, China), and aqueous DNA was pipetted from just above the Chelex^**®**^ resin, ensuring the resin remained in the tube.

PCR reactions were undertaken using a 1/10 dilution of each DNA extraction, amplifying the samples using immolase polymerase and the “universal” arthropod primer pair LCO1490/HCO2198 (Folmer et al. 1994). Reactions contained a final concentration of 1x Standard Taq Reaction Buffer (New England Biolabs, Massachusetts, USA), 2.5 mM MgCl2, 0.5 μM each primer, 0.2 mM dNTPs, 2.4 U IMMOLASE DNA Polymerase (Bioline, London, UK) and 3 μL diluted DNA, in a reaction volume of 30 μL. Amplicons were sent to the Australian Genome Research Facility (AGRF) for sequencing, before forward and reverse sequences were assembled and trimmed using Geneious version 9.1.8 (https://www.geneious.com). Sequences were searched against the Genbank database (http://www.ncbi.nlm.nih.gov) and cross referenced with the Barcode of Life Data System database (BOLD; http://www.barcodinglife.org; (Ratnasingham & Hebert 2007) to ascertain their identity.

### 2.5. Online databases

Online databases were used to collect records of aphidiines up to and including June 2020. Any uncertain morphological identification was recorded as such, as specimens were unable to be morphologically identified based on photographs alone. Data were acquired from the online database, the Atlas of Living Australia (ALA; http://www.ala.org.au) and the Global Biodiversity Information Facility (GBIF; https://www.gbif.org). Host-parasitoid associations could not be ascertained from these datasets; these databases were used only for distribution mapping of aphidiine species.

### 2.6. Distribution mapping & host associations

Species distribution maps were created for each species from the repeat sampling, opportunistic sampling, insect depository, and online collection datasets. Host association matrices and food webs were created in R version 4.0.1 (R Core Team 2020), using RStudio version 1.3.959 (RStudio Team 2020) and the bipartite package (Dormann 2020).

## 3. Results & Discussion

### 3.1. Aphidiine diversity

In total, 2973 aphidiines were collected during the repeat sampling between 2017 and 2019, and 2009 aphidiines through the opportunistic sampling between 2016 and 2019. 487 aphidiines were collated from the historic insect collections and 56 from the online databases. In total, these 5525 specimens constituted a total of 23 species and seven genera. Across all samples, *D. rapae* was the most common aphidiine at 72.05%, followed by *Aphidius ervi* at 6.05%, *Aphidius colemani* at 5.85% and *Lysiphlebus testaceipes* at 5.30% (Fig. 3). Across the states and territories, 1034 aphidiines were sampled and collated from NSW, 60 from QLD, 1015 from SA, 369 from TAS, 2231 from VIC and 640 from WA. A further 176 aphidiines were collated from Australian Capital Territory (ACT). There were no records or collections from the Northern Territory (Fig. 4).

**Figure 3:**
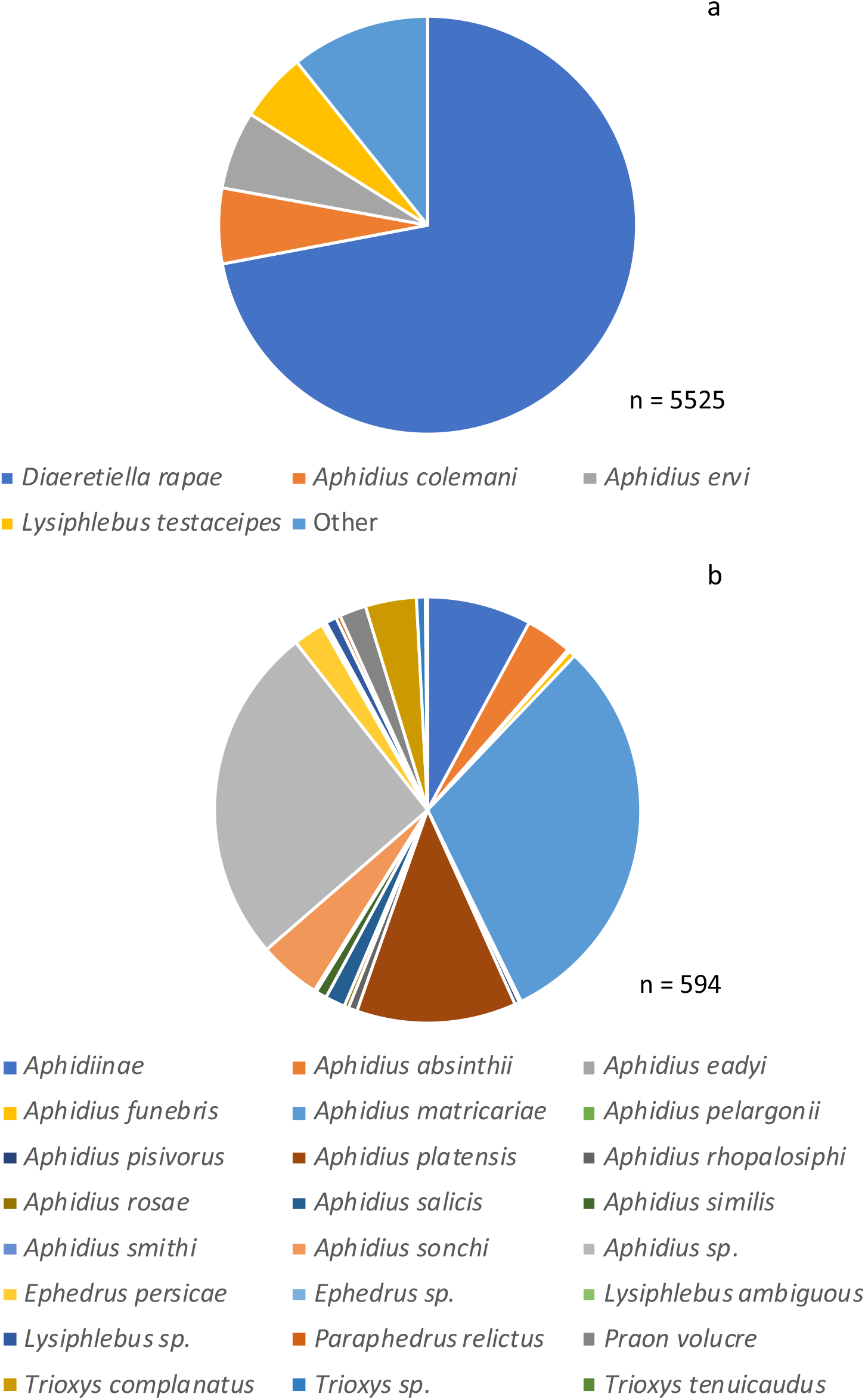
Species composition of aphidiines from sampling and collated data, with a) showing the common species and b) showing the breakdown of the ‘other’ aphidiines category.

**Figure 4:**
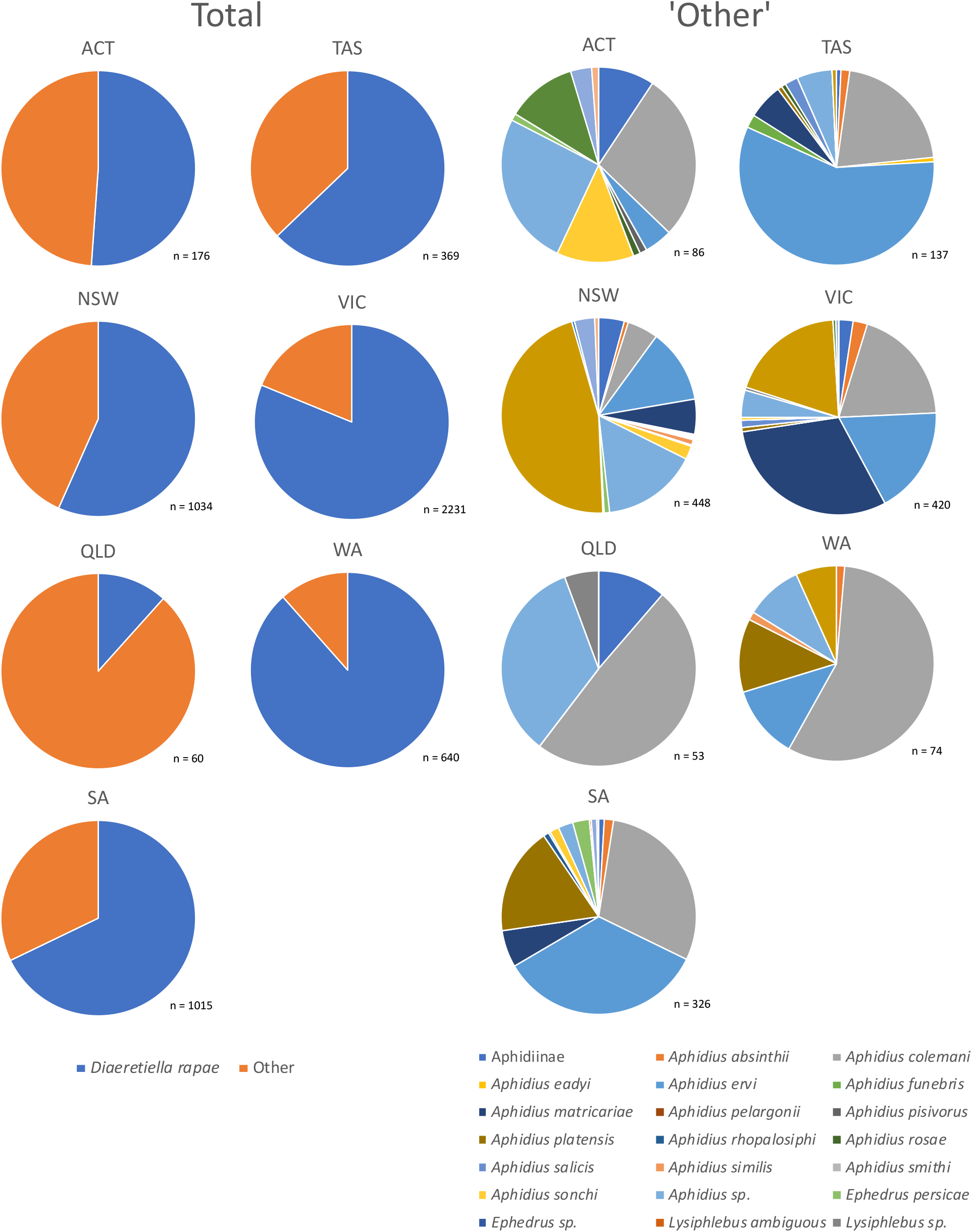
Species composition of aphidiines for each state and territory from sampling and collating data [ACT = Australian Capital Territory, NSW = New South Wales, QLD = Queensland, SA = South Australia, TAS = Tasmania, VIC = Victoria, WA = Western Australia].

### 3.2. Distribution mapping

The distribution of aphidiines across Australia was mapped using all sampled and collated data (Fig. 5). Aphidiine species found at ten or more sampling points were plotted separately, with Fig. 6 covering the most commonly recorded species and the remainder plotted in Fig. S1. The lack of data from the Northern Territory is largely explained by the lack of grain and oilseed crops grown there. The southern grain belt extends from WA across the bottom of the country through VIC and into NSW. QLD records tended to be from horticultural crops, although the lower region of this state is suitable for grain growing. Further research into aphidiines within the NT and northern QLD would be informative. Aphidiine presence generally correlates with the southern grain belt, likely due to the abundance of preferred host plants and climate and increased sampling in that region. Records of *D. rapae* span a wider range than for the other species, reaching a more northerly point (Fig. 6). *Aphidius colemani* and *A. ervi* are only found on the eastern side of the Spencer Gulf in SA, like all other aphidiine species shown in Fig. S1. *Diaeretiella rapae* was the only species recorded or collected on the Eyre Peninsula of SA. *Aphidius ervi* is common in TAS, and does not reach QLD, unlike *D. rapae* and A*. colemani*, which constituted half of the species recorded in that state (Figs. 4, 6).

**Figure 5:**
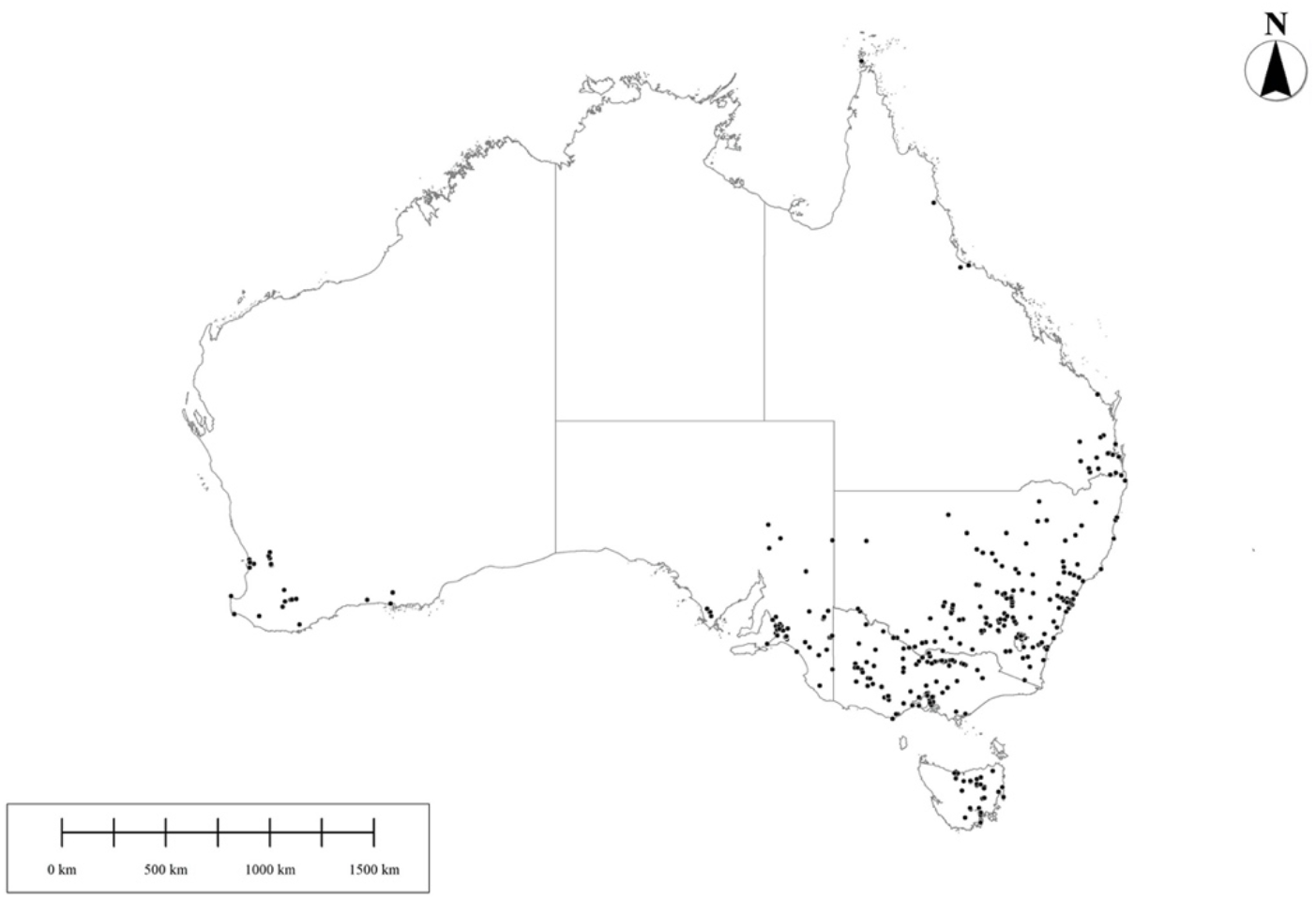
Aphidiine distribution within Australia, based on all collated data.

**Figure 6:**
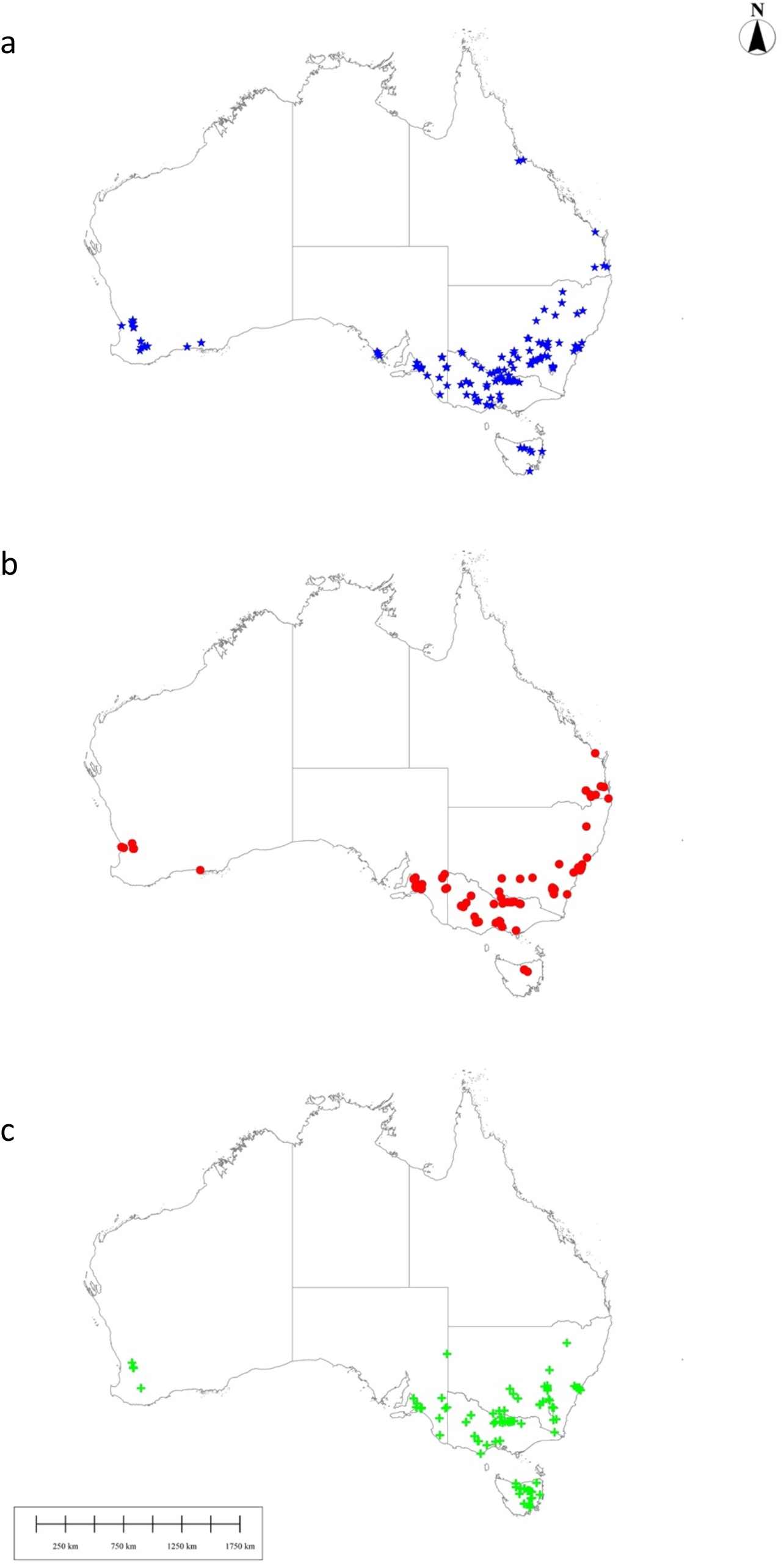
Distribution of a) *D. rapae*, b) *A. colemani*, and c) *A. ervi* in Australia, based on all collated data.

### 3.3. Taxonomy of sampled aphidiines

Field collections of aphidiines (collected both through repeat and opportunistic sampling) were identified to species level. Of the 4982 aphidiines identified, 3488 were reared from collected mummies, 108 were pan trapped, 170 were sampled by sweep netting, 747 were vacuum sampled, and 469 were directly sampled.

The key below is based on the eleven species of aphidiine that we sampled within grain production landscapes between 2016 and 2019.

#### 3.3.1. Dichotomous key to aphidiines in Australian grain production landscapes

Key to females of Aphidiinae species from Australia (adapted from Rakhshani et al. (2015)), (with notes on males, when present).

1. RS+M vein present in fore wing (Fig. 7) Notaulices well developed and distinct……..………….…………. ***Praon volucre* (Haliday, 1833)**

-RS+M vein absent (Fig. 7). Notaulices incomplete or absent………………………………..……………. **2**

**Figure 7:**
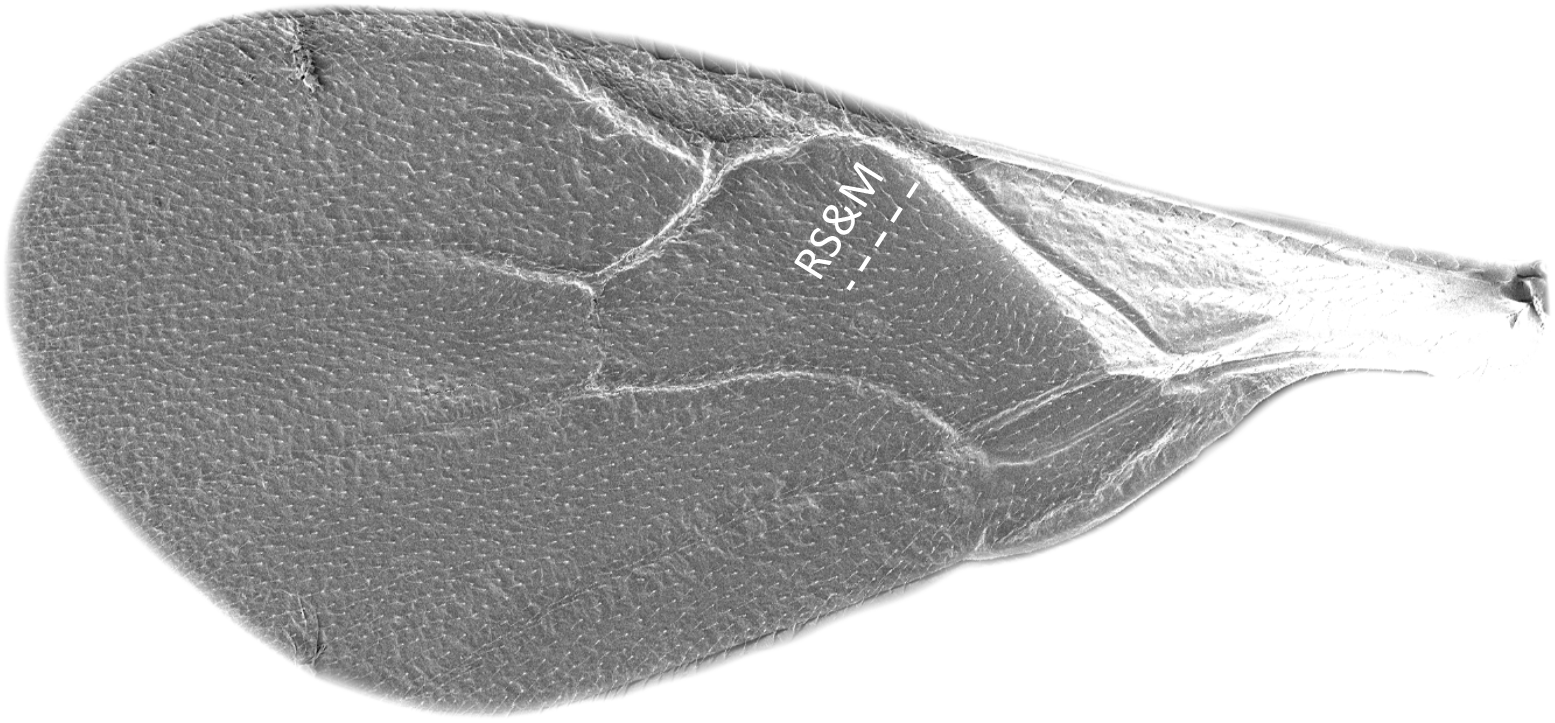
Fore wing of *Aphidius sonchi* [Position of lacking RS&M vein illustrated].

2. Fore wing M&m-cu vein incomplete (Fig. 8a) or absent (Fig. 8b) ..………….…………….……… **10**

**Figure 8:**
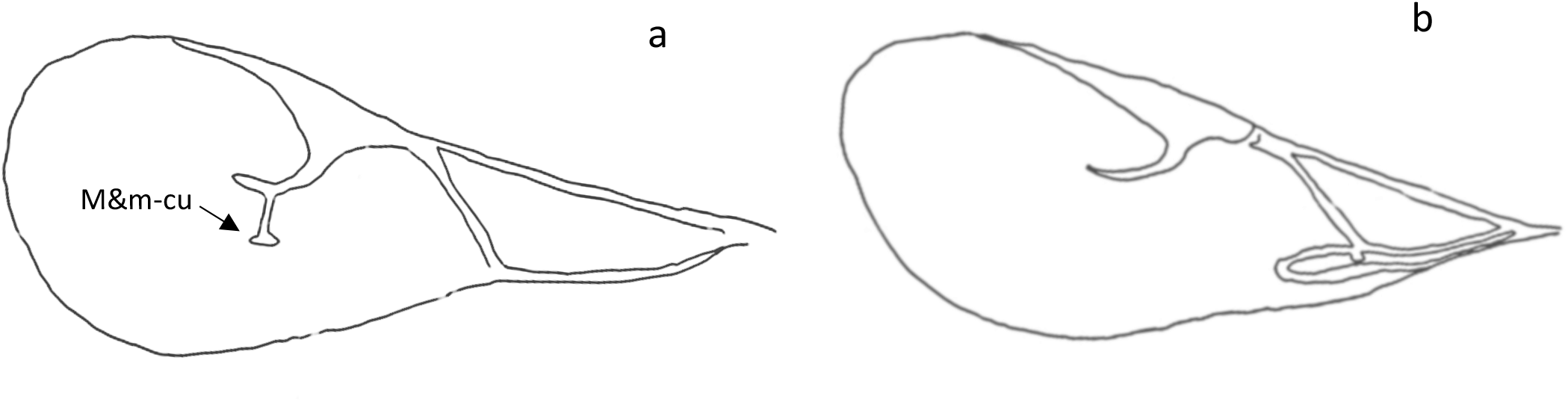
Fore wing of a) *Lysiphlebus testaceipes*, b) *Diaeretiella rapae*.

-Fore wing M&m-cu vein present and complete (Fig. 9) ……..………….………………………………….. **3**

**Figure 9:**
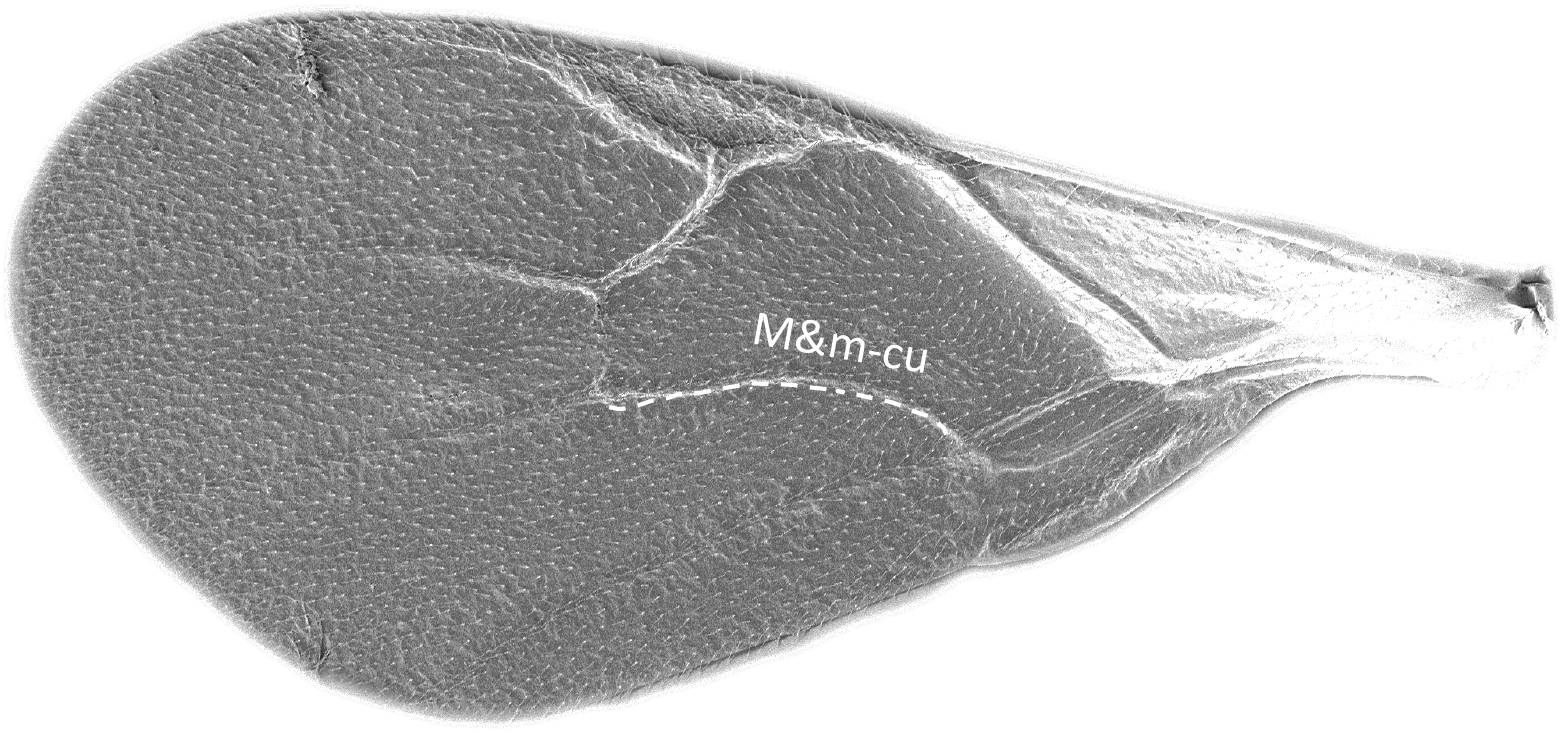
Fore wing of *Aphidius sonchi*.

3. Anterolateral area of petiole rugose (Fig. 10) ..………….……….. ***Aphidius ervi*, Haliday, 1834**

(Antennae 17-19-segmented in females, 19-21-segmented in males)

**Figure 10:**
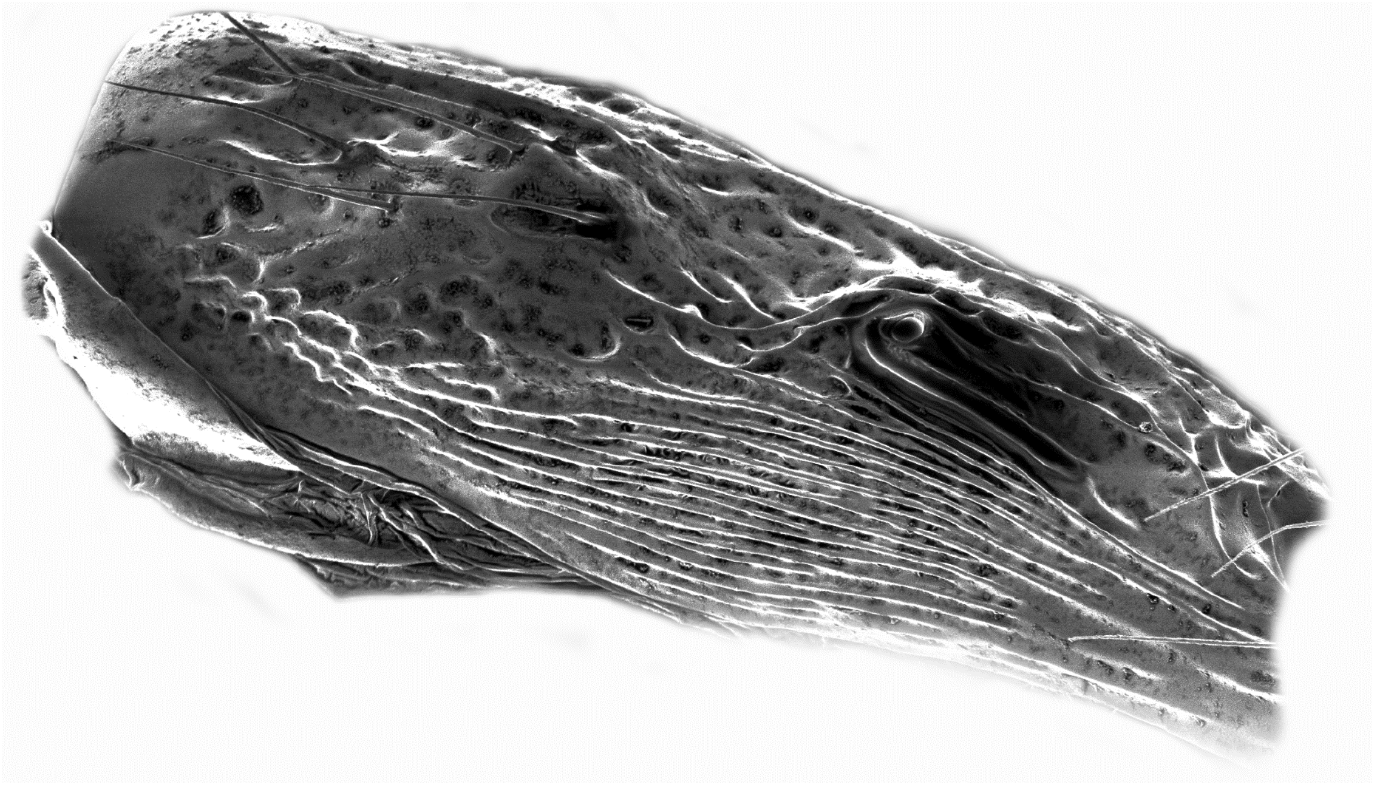
Anterolateral area of petiole of *Aphidius ervi*.

-Anterolateral area of petiole costate/costulate (Figs. 11a & b) .…………………..………….………… **4**

**Figure 11:**
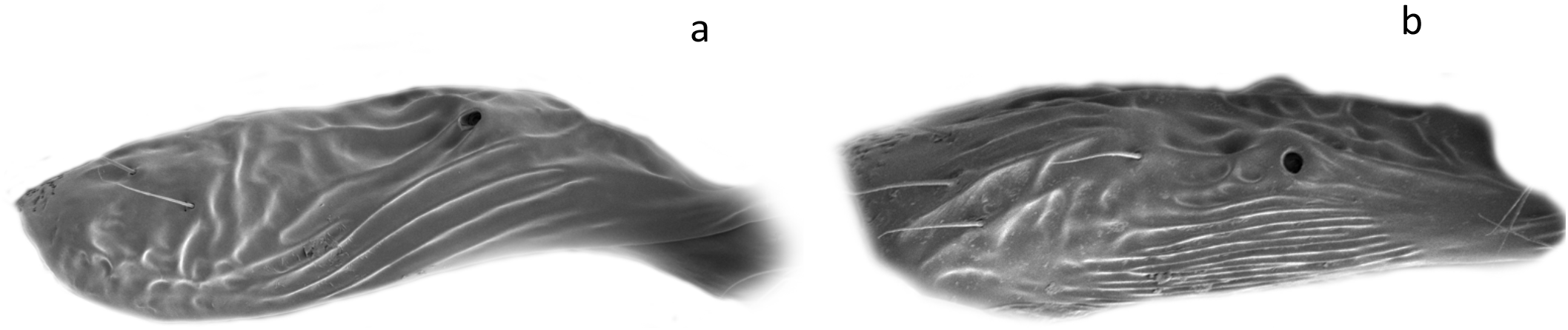
Anterolateral area of petiole of a) *Aphidius colemani*, b) *Aphidius matricariae*.

4. Anterolateral area of petiole costate (Fig. 11a) ……………..…………………………………….…….….. **5**

-Anterolateral area of petiole costulate (Fig. 11b) ……..……………….……………….…….………..…….. **6**

5. Fore wing R1 vein almost equal to stigma length, anterolateral area of the petiole with ~3 distinct costae (Fig. 12a) …………………………………………………. ***Aphidius colemani*, Viereck, 1912**

(Antennae 13-16-segmented in females, 16-19-segmented in males)

**Figure 12:**
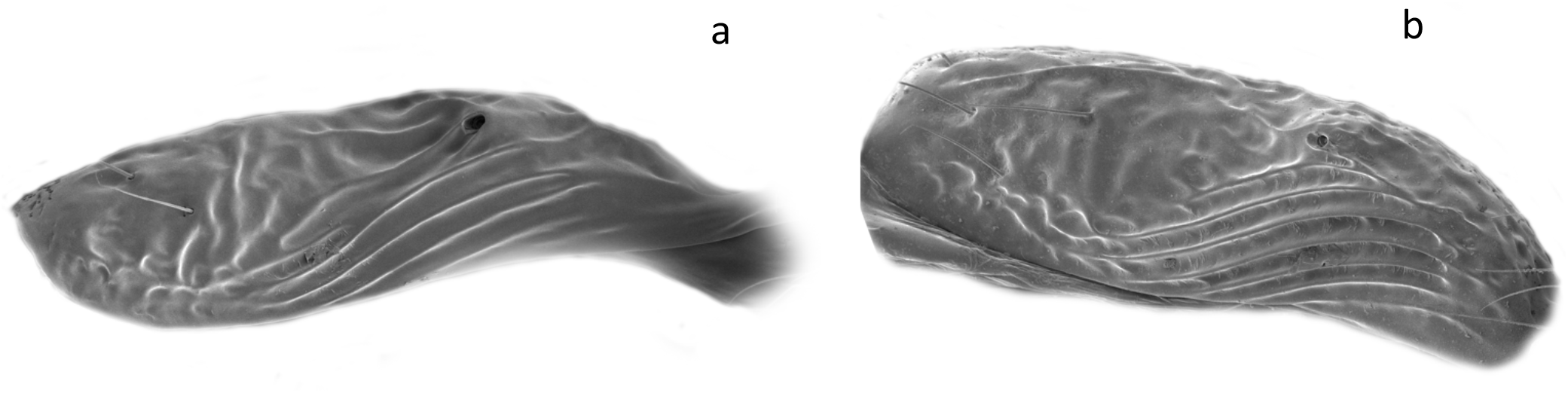
Anterolateral area of petiole of a) *Aphidius colemani*, b) *Aphidius platensis*.

- Fore wing R1 vein about one-third shorter than the stigma length, anterolateral area with 5-7 costae (Fig. 12b) ………………………………………………………… ***Aphidius platensis*, Brèthes, 1913**

(Antennae 15-segmented in females, 18-19 segmented in males)

6. Labial palpi 3-segmented (Fig. 13a) .…………………….…..……………………………………..…….…….… **7**

- Labial palpi ≤ 2-segmented (Fig. 13b) .…………………………………………………………….…..……… **9**

**Figure 13:**
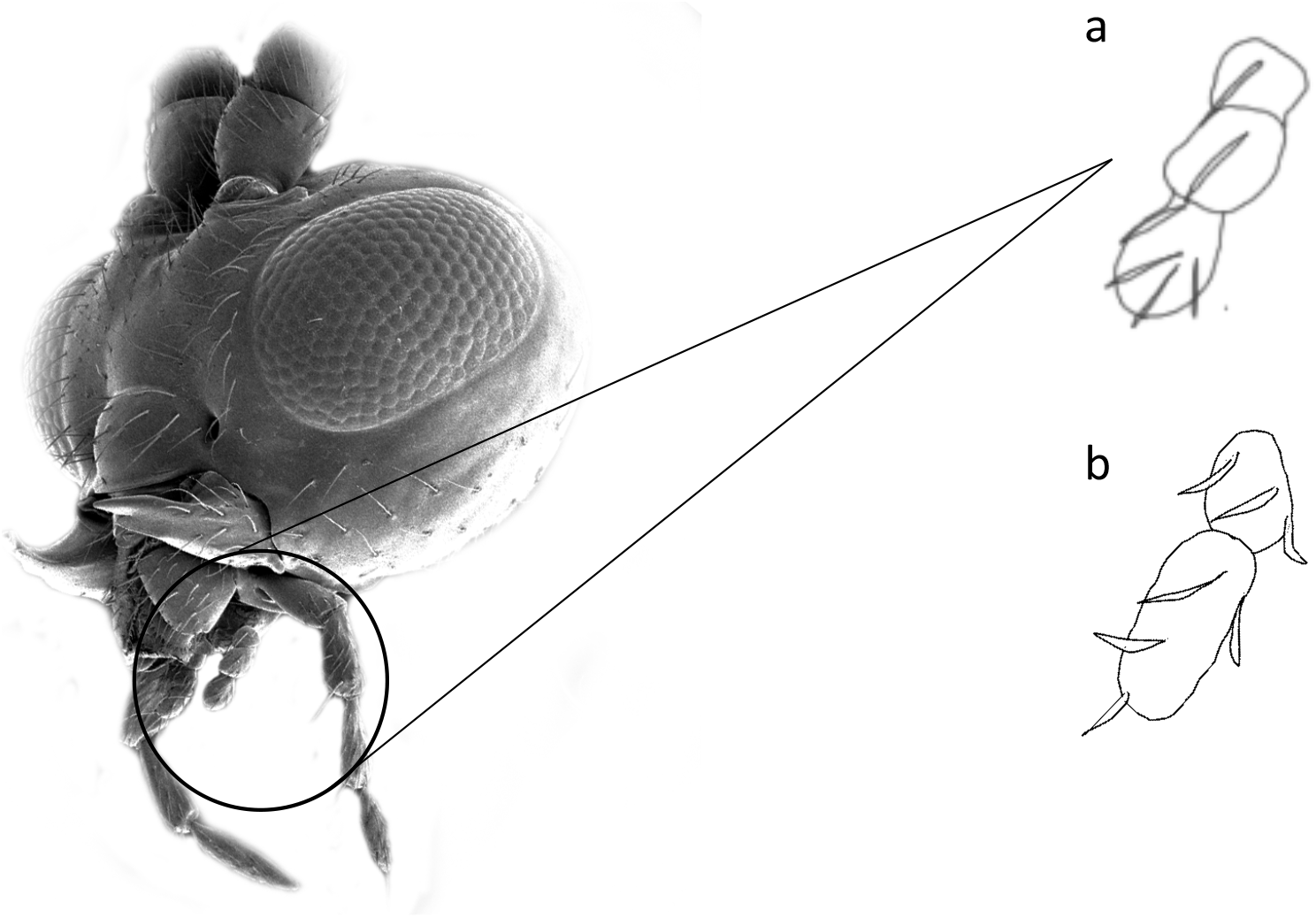
Labial palpi of a) *Aphidius ervi*, b) *Aphidius matricariae*.

7. Ovipositor sheath strongly prominent and elongate ………………………………………………………… ……………………………………………………………………………………… ***Aphidius funebris*, Mackauer, 1961**

(Antennae 17-19 segmented in females, 20-21 segmented in males)

- Ovipositor sheath broad (Fig. S2a)……………………………………………………………………………………..**8**

8. Maxillary palps with four palpomeres …….. ***Aphidius rhopalosiphi*, De Stefani Perez, 1902**

(Ovipositor sheath strongly concave in its anterior-dorsal margin, widening towards base. Petiole with unclear central carina. Antennae 16-17-segmented in females, 18-segmented in males)

- Maxillary palps with three palpomeres……………..…………….. ***Aphidius sonchi*, Marshall, 1897**

(Antennae 15-16-segmented in females, 18-segmented in males)

9. Antennae 14-15-segmented ………………..……..………… ***Aphidius matricariae*, (Haliday, 1834)**

(Antennae 17-18-segmented in males)

- Antennae 16-17-segmented….……………………………………. ***Aphidius absinthii*, Marshall, 1896**

(Antennae 17-19-segmented in males)

10. Fore wing M&m-cu vein incomplete, reduced in anterior part (Fig. 8a). Propodeum smooth (Fig. 14a) ………..…………….…………………..……… ***Lysiphlebus testaceipes* (Cresson, 1880)**

(Antennae 12-15-segmented in females, 13-15-segmented in males)

**Figure 14:**
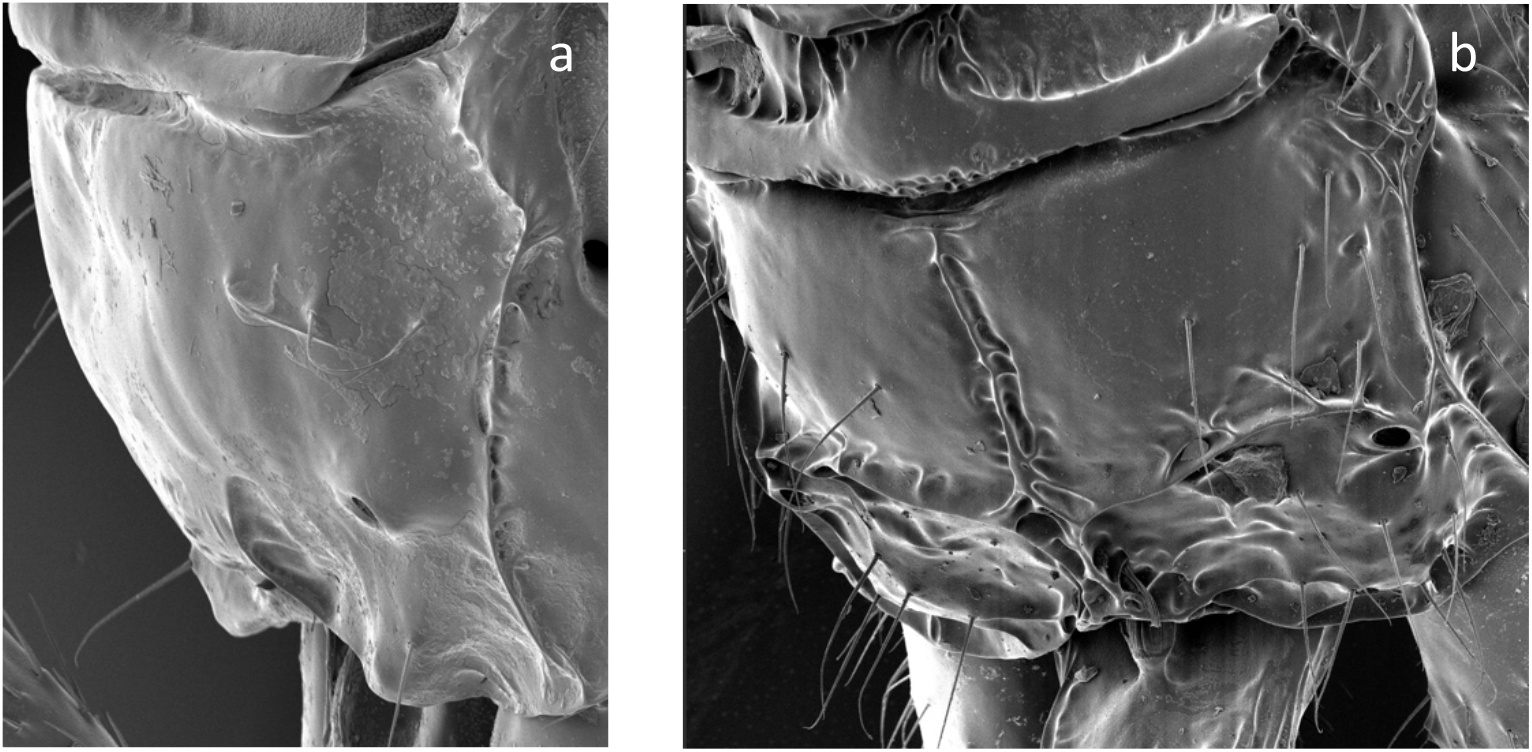
Propodeum of a) *Lysiphlebus testaceipes*, b) *Diaeretiella rapae*.

-Fore wing M&m-cu vein completely absent (Fig. 8b). Propodeum carinated (Fig. 14b) ………… ……………………………….…………………………………………………… ***Diaeretiella rapae* (M’Intosh, 1855)**

(Hypopygium without prongs. Fore wing stigma elongate triangular. Propodeum with narrow pentagonal areola. Ovipositor sheath short, straight, apically truncated. (Antennae 13-14 segmented in females, 16-17 segmented in males)

[Notes on sexing can be found in the supplementary material (Fig. S3)].

### 3.4. Host-parasitoid associations of sampled aphidiines

Of those parasitoids reared from aphids collected in grain production landscapes between 2016 and 2019, *D. rapae* was found to be the most dominant species (reflecting 73.54% of all parasitoids), followed by *Lysiphlebus testaceipes* (Cresson) (7.63%), *A. ervi* (6.57%), and *A. colemani* (5.93%). Other parasitoid species each constituted <5% of the total wasps reared (Fig. 15).

**Figure 15:**
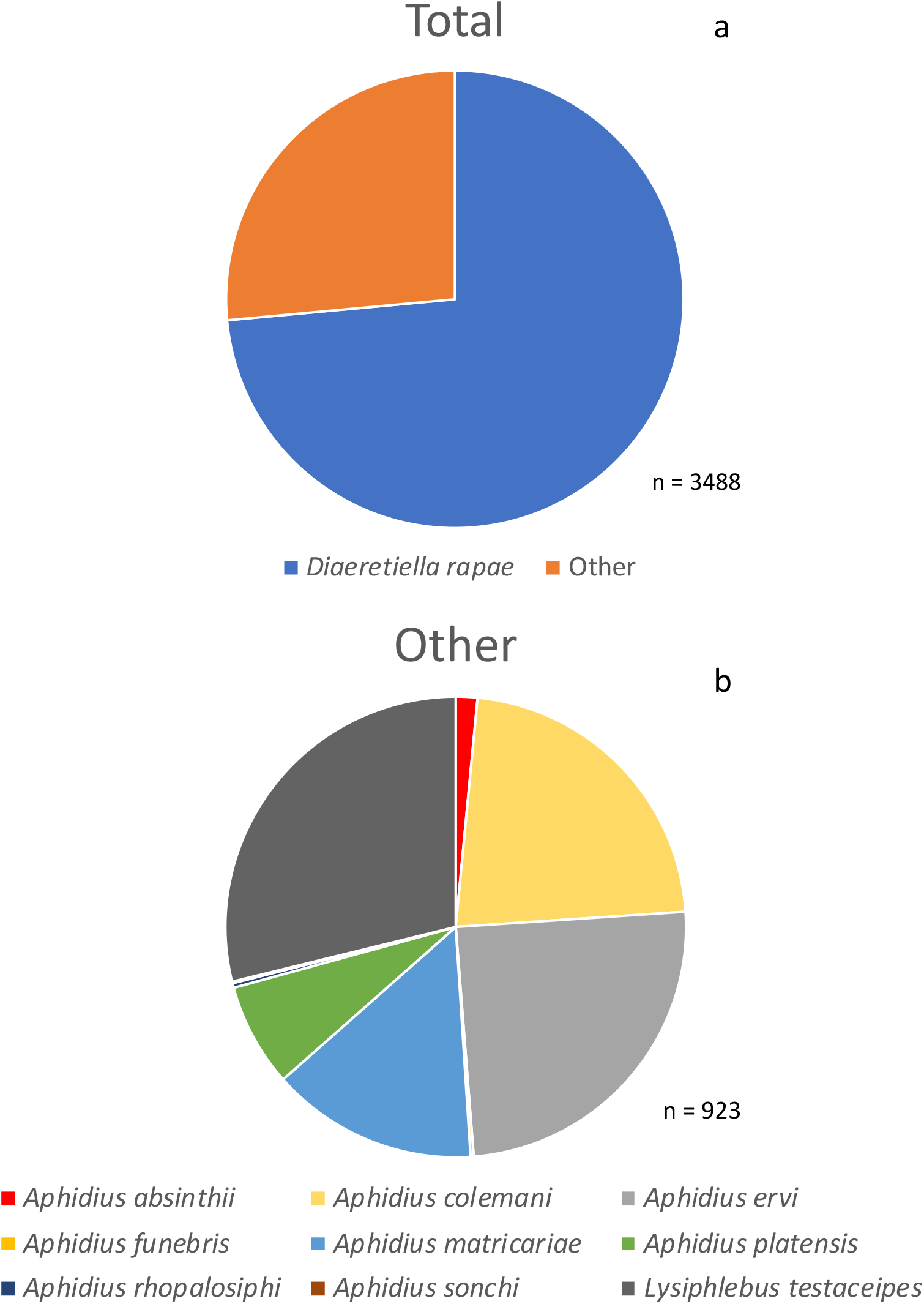
Species composition of aphidiines reared from aphids, with a) showing the total and b) showing the breakdown of the ‘other’ aphidiines category.

Across Australia, aphid host-aphidiine associations were analysed from wasps reared from aphids collected during the repeat and opportunistic sampling trips (Fig. 16). *Diaeretiella rapae* dominated brassica aphid (*M. persicae*, *B. brassicae* and *L. erysimi*) parasitism, yet *L. testaceipes* dominated *R. maidis* and *D. noxia* parasitism. *Aphidius funebris* was the only aphidiine found parasitising *A. pisum*. *Aphidius ervi* dominated *M. dirhodum* parasitism, and *A. matricariae* dominated *R. padi* parasitism*. Myzus persicae* was parasitised by the greatest number of aphidiine species, which could be due to its polyphagous nature and ability to feed on over 40 plant families (Blackman & Eastop 2000; 2007). *Aphidius platensis* was only found parasitising brassica aphids, suggesting the host plant may be a greater attractant for this parasitoid species than the aphid host, as volatiles are known to play a role in aphidiine foraging behaviour, although this can vary depending on the aphidiine species (Müller & Godfray 1999).

**Figure 16:**
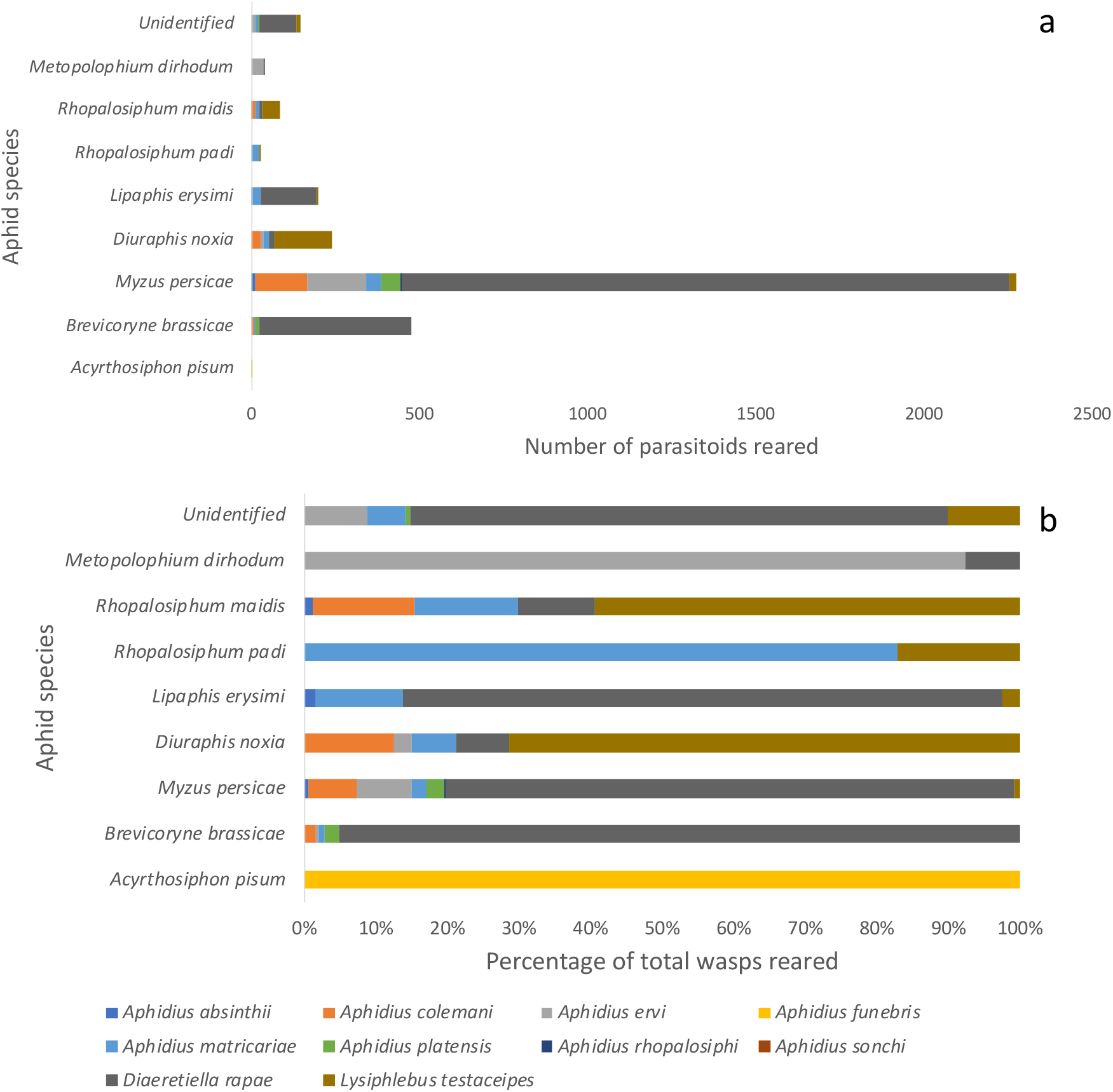
Composition of parasitoids reared from aphid species in grain production landscapes within Australia as a) raw numbers, and b) proportions of total.

Aphid host-aphidiine associations were analysed by state (Fig. S4). In all states, when rearing from brassica aphids, *D. rapae* was the dominant wasp species, apart from in QLD. Although numbers were low, only *A. colemani* was reared from *M. persicae* in QLD. This is likely because the host plant was broccoli (*Brassica oleracea* L.). Although *D. rapae* is often reared from aphids on broccoli, *A. colemani* is recorded as being conditioned to oviposit within the plant it has developed on, rarely switching between host species in the field (Coll & Hopper 2001). QLD temperatures and humidity are usually higher than for the other states in which mummies were sampled. This could contribute to the relatively higher abundance of *A. colemani* in QLD as there is a correlation between temperature and population density of *A. colemani*, which is not the case for *D. rapae* (Saleh et al. 2006).

*Diuraphis noxia* reared mostly *L. testaceipes* in NSW, however it reared mostly *A. colemani* in TAS. Again, this could be attributed to the host conditioning of *A. colemani*, and variation of host plants from which the *D. noxia* mummies were collected, which covered a wide range of host plants (Table S1). *Lysiphlebus testaceipes* also has a lower tolerance than *A. colemani* to cold temperatures (Jones et al. 2003), and so is likely to be outcompeted in the cooler climate of Tasmania during spring, when the majority of samples were taken in this state.

Plots of tri-trophic host associations show that many of the parasitoids were reared from a greater range of host plants than host aphids (Table 2; Fig. S5). *Aphidius ervi* had the broadest plant host range, followed by *D. rapae*, while *A. matricariae* and *D. rapae* had the greatest range of aphid hosts. Žikić et al. (2017) investigated the host range patterning of Aphidiinae and found *D. rapae* to have the greatest range of associated hosts (between 12 and 168). However, international literature suggests that the majority of parasitoids are found in association with 1-2 host aphid species (Nazari et al. 2012), which is far fewer than observed for most parasitoids characterized here.

**Table 2:** Host range of aphids and plants for each aphidiine species reared

| Aphidiine species | Number of host aphids attacked | Number of host plants attacked |
| --- | --- | --- |
| *Aphidius absinthii* | 5 | 3 |
| *Aphidius colemani* | 2 | 7 |
| *Aphidius ervi* | 4 | 10 |
| *Aphidius funebris* | 1 | 1 |
| *Aphidius matricariae* | 6 | 7 |
| *Aphidius platensis* | 2 | 3 |
| *Aphidius rhopalosiphi* | 1 | 1 |
| *Aphidius sonchi* | 1 | 1 |
| *Diaeretiella rapae* | 6 | 9 |
| *Lysiphlebus testaceipes* | 5 | 7 |

Apart from those parasitoids reared from only one host aphid, all aphidiine species were reared from both brassica and cereal aphids. Due to the higher proportion of canola paddocks sampled during both the random and opportunistic sampling, and the dominance of *M. persicae* within this crop, most trophic interactions occurred between this aphid species and the dominant aphid parasitoid, *D. rapae* (Fig. 17). Similarly, in France between 1998 and 2001, *D. rapae* was one of the two major aphidiines (the other being *A. matricariae*) found in canola parasitising *M. persicae* (Desneux et al. 2006). Conversely, in India, *D. rapae* was found to parasitise *B. brassicae* more commonly than other aphids within canola, followed by *L. erysimi* and *M. persicae* (Desh & Lakhanpal 1998)*. Diaeretiella rapae* was the dominant aphidiine species within canola paddocks in central Oklahoma, however parasitism of *M. persicae* tended to be lower than for *L. pseudobrassicae* (Davis) (Elliott et al. 2014). This suggests tri-trophic interactions that have been established overseas may have little resemblance to those that have evolved in Australia.

**Figure 17:**
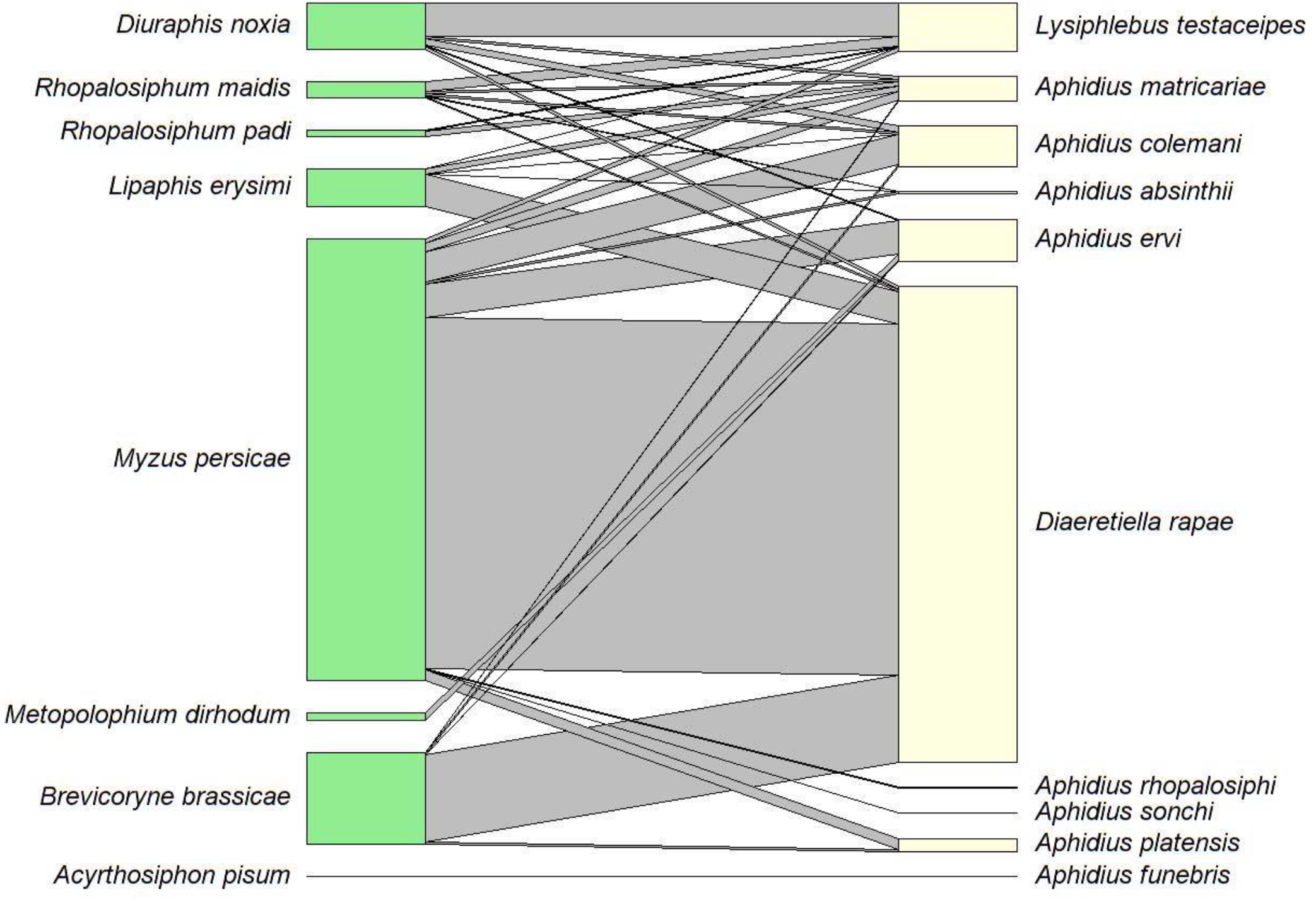
Quantitative food web of host associations between aphids and aphidiines, for each aphidiine species reared during the random and opportunistic sampling [box sizes indicate number of records in relation to other species].

### 3.5. Conclusions

Preliminary studies undertaken by Carver and Starý (1974) suggested the aphidiid fauna within Australia was smaller than expected, however they stated that more records were required. A subsequent study by Carver (1992) found only 15 species of aphidiine within Australia. It is surprising, almost 30 years later, that limited diversity of this parasitoid group remains the case in grain production landscapes, with even fewer species recorded during this study. *Myzus persicae*, the main aphid host, is recorded as being widespread globally due to human-mediated transport (Margaritopoulos et al. 2009), and there has been concern over the spread of aphids through international airline travel and horticultural cargo (Venette & Ragsdale 2004). The parasitoids associated with these aphids and others might be expected to be transported in the form of developing larvae within mummies. For example, Braconidae and Ichneumonidae have been found within litter and soil transported from the North Island to the South Island of New Zealand (McNeill et al. 2006). It is therefore unexpected that the number of aphidiine species remains low.

Globally, there are more than 400 species of Aphidiinae, from over 50 genera (Mackauer & Starý 1967; Starý 1988). This subfamily is important in the biological control of crop pests, yet can be found within numerous habitats, including in wetlands (Tomanović et al. 2012), on ornamental plants (Kavallieratos et al. 2013), and in forests (Kaliuzhna 2019). A survey in India found 125 species of aphidiine across 22 genera over numerous host plants and aphids (Akhtar et al. 2011). Forty species of aphidiine were reared from 50 species of aphid in Slovenia between 2006 and 2010 (Kos et al. 2012), while 81 species (from 19 genera) of aphidiine were recorded parasitizing over 200 species of aphid in the USA (Pike et al. 2000). Rakhshani et al. (2015) found 16 aphidiine species in Malta. Within Iran, Rakhshani et al. (2008a) identified 17 species of *Aphidius* between 2001 and 2005. Olmez and Ulusoy (2003) found 16 species of aphid parasitoids between 1998 and 2000 within the Diyarbakir Province, Turkey. Another study in Turkey, in Kahramanmaras, found 18 aphidiine species on 30 hosts (Aslan et al. 2004). Perhaps the low local diversity compared to the diversity found in these overseas studies reflects the dominance of *D. rapae* which may be restricting the establishment of other aphidiines, particularly within brassica crops. However, there is clearly a need to investigate aphidiine diversity further in Australia, particularly in other cropping systems, in horticultural settings, and in native vegetation, including those in northern Australia.

Although there are a range of aphidiine species present within Australian grain production landscapes, *D. rapae* dominates across Australia, covering approximately 75% of all parasitoids recorded. This is not always the case at a local scale, however, with host plants likely contributing to the enhancement of aphidiine parasitism either by particular species in certain geographic regions or on certain aphid hosts. The aphidiine fauna in Australia is similar to that found elsewhere in the world, with overlap of species described in other studies, such as Kos et al. (2012). No new species were recorded in our surveys. The tritrophic interactions we describe may help to inform aphid management in grains landscapes, with many aphidiines able to host swap across aphid species and host plant species. IPM strategies that include aphidiines could promote these parasitoids on pest and non-pest aphids utilising plants adjacent to crops.

## Acknowledgements

We would like to acknowledge the growers and agronomists who assisted with this project and provided access to their land, in addition to those people who collected specimens on our behalf (Elia Pirtle, Dusty Severtson, Amber Balfor-Cunningham, Matthew Binns, Jo Holloway, Rachel Wood, Caitlin Langley, Maarten van Helden, Tom Heddle, Xuan Cheng, and others). We thank those individuals who provided specimens from insect collections, and those who assisted with the organisation of such material viewing. We would like to thank Sarina Macfadyen for her supervision. Electron microscopy and training was carried out at the Bio21 Advanced microscopy facility, under the supervision of Roger Curtain, who we would like to thank for his assistance. We further extend our gratitude to Melanie Carew, Vanessa White, Xuefen Xu, Moshe Jasper, and Katie Robinson, for their assistance with genetic work and training, Erica Marshall with her assistance in R programming, Ehsan Rakhshani for his words of wisdom, and to Vincent Chea for his GIS expertise. This research was supported by a GRDC investment that seeks to deliver new knowledge to improve the timing of pest management decisions in grain crops to grain growers: CSE00059. This project was undertaken by the Commonwealth Scientific and Industrial Research Organization, in partnership with Cesar Australia, NSW Department of Primary Industries, the South Australian Research & Development Institute, the University of Melbourne, and the Western Australian Department of Primary Industries and Regional Development. This work was further supported by the Michael Mavrogordato award from the Australian Native Animal Trust, the Albert Shimmins Fund, and the Australian Grains Pest Innovation Program.

