## Supplementary materials for "Taxonomy, distribution and host relationships of aphidiine wasps (Hymenoptera: Aphidiidae) parasitizing aphids (Hemiptera: Aphididae) in Australian grain production landscapes"

*Table S1:* Summary of host plants from which parasitoids were reared from aphid mummies across all field surveys between 2016 and 2019.

| | **HOST PLANT** | | | | | | | | | | | | | | | |
|---|---|---|---|---|---|---|---|---|---|---|---|---|---|---|---|---|
| ***STATE OR TERRITORY*** | *Brassica napus* | *Malva sp.* | *Triticum aestivum* | *Hordeum vulgare* | *Brassica oleracea var. italica* | *Raphanus raphanistrum subsp. sativus* | *Phalaris sp./Bromus sp.* | *Sonchus oleraceus* | *Arctotheca calendula* | *Prunus persica* | *Avena sativa* | *Pisum sativum* | *Hordeum sp.* | *Brassica oleracea var. capitata* | *Brassica rapa subsp. chinensis* | *Phaseolus vulgaris* |
| **ACT** | X | | | | | | | | | | | | | | | |
| **NSW** | X | | X | X | | X | X | | | | | | X | X | | |
| **QLD** | | | | | X | | | | | | | | | | | |
| **SA** | X | | | | | | | | | | | | | | | X |
| **TAS** | X | | X | X | | X | X | X | X | | X | X | | | | |
| **VIC** | X | X | X | X | X | X | | | | X | | | X | X | X | |
| **WA** | X | | | | | | | | | | | | | | | |

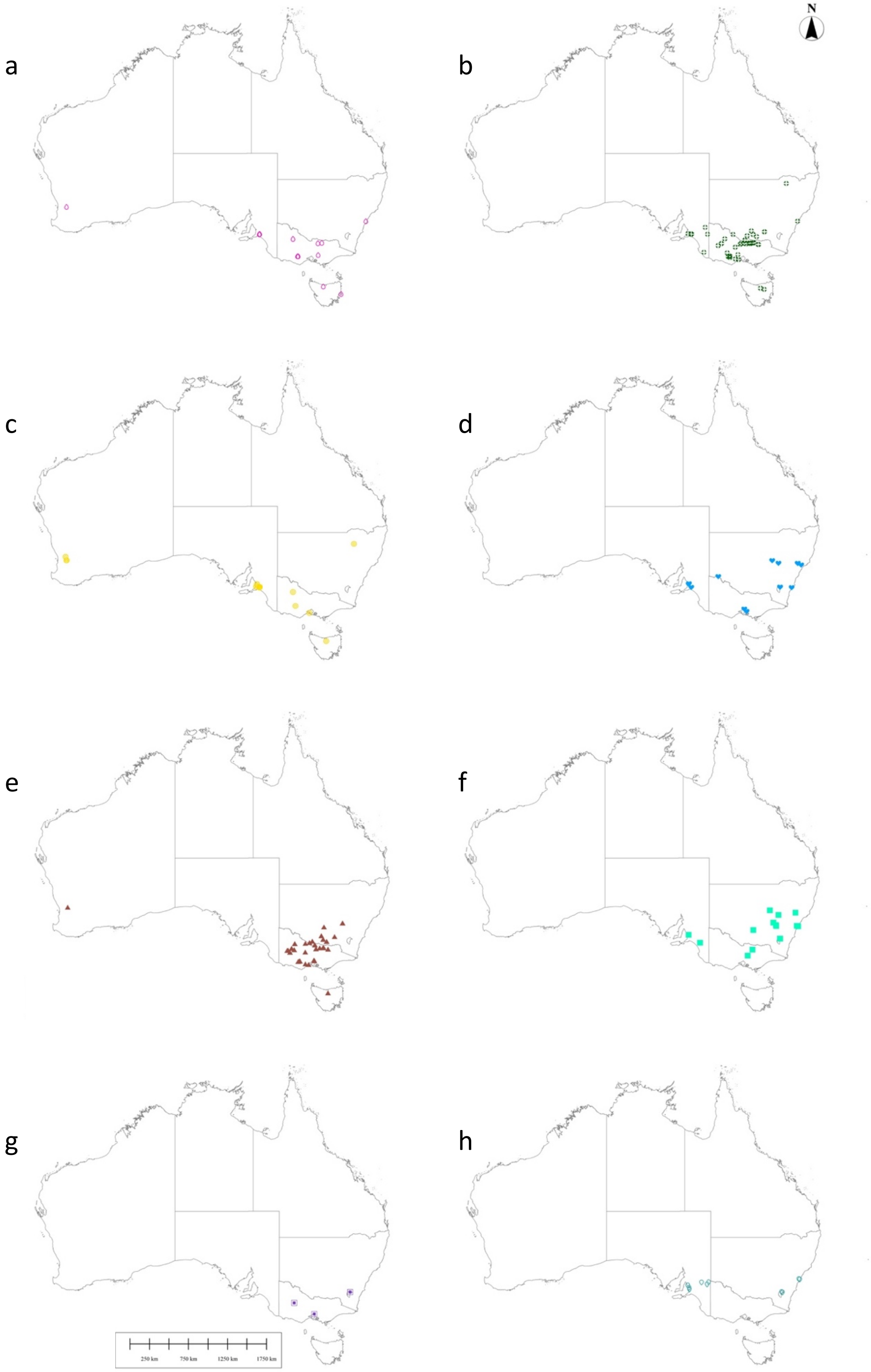


*Fig. S1:* Aphidiine distribution across Australia, based on all collated data for a) *Aphidius absinthii*, b) *Aphidius matricariae*, c) *Aphidius platensis*, d) *Aphidius sonchi*, e) *Lysiphlebus testaceipes,* f) *Trioxys complanatus,* g) *Praon volucre,* and h) *Ephedrus persicae.*

*Note on sexing*

Sexing can be undertaken from external genital structures, with females possessing an ovipositor sheath (3rd valvula), which can vary in shape, depending on the species (Fig. S2a), and males possessing distinguishable gonoforceps, paired lateral parts of the male genitalia (Fig. S2b). Additionally, the number of visible metasomal sternites within males and females differ, with five visible in females (Fig. S3a) and seven visible in males (Fig. S3b), (prior to the hypopygium).

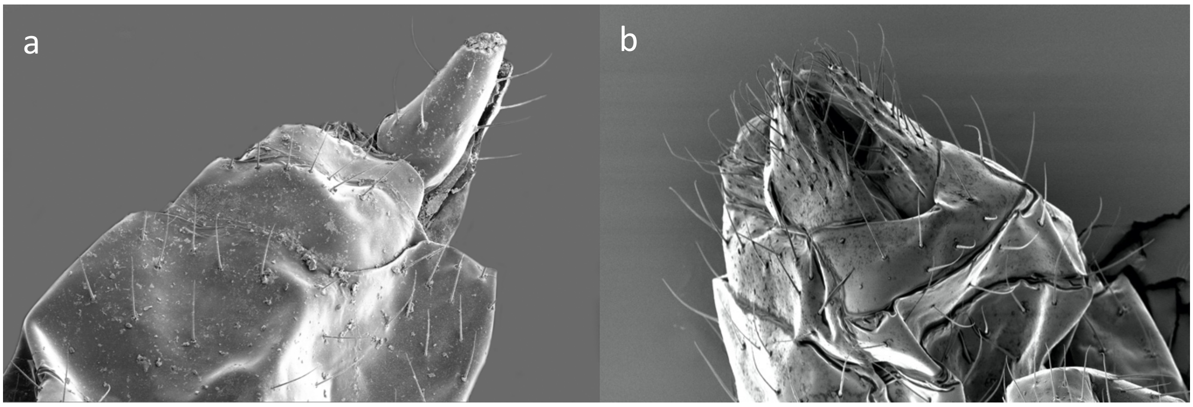

*Figure S2*: Genitalia of a) *Diaeretiella rapae* female, b) *Aphidius ervi* male.

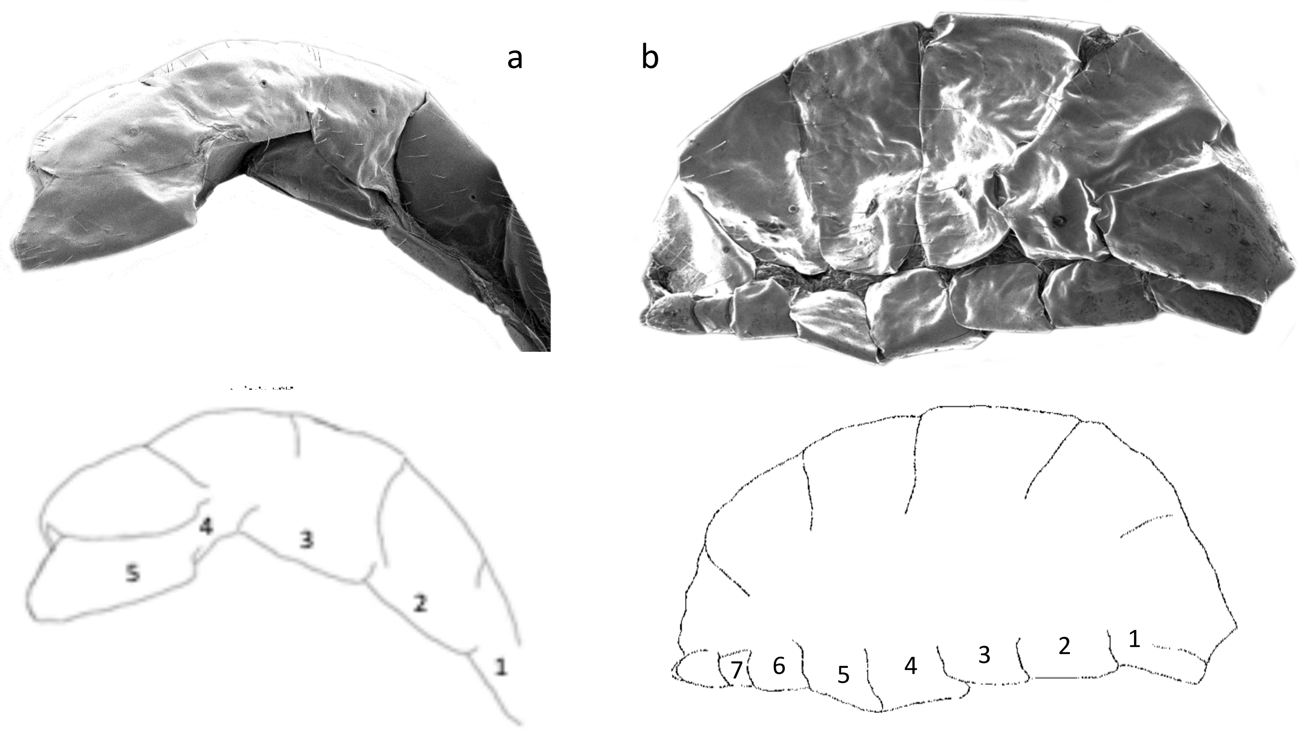

*Figure S3*: Sternites of a) a female, and b) a male [with respective diagrams illustrating number of sternites].

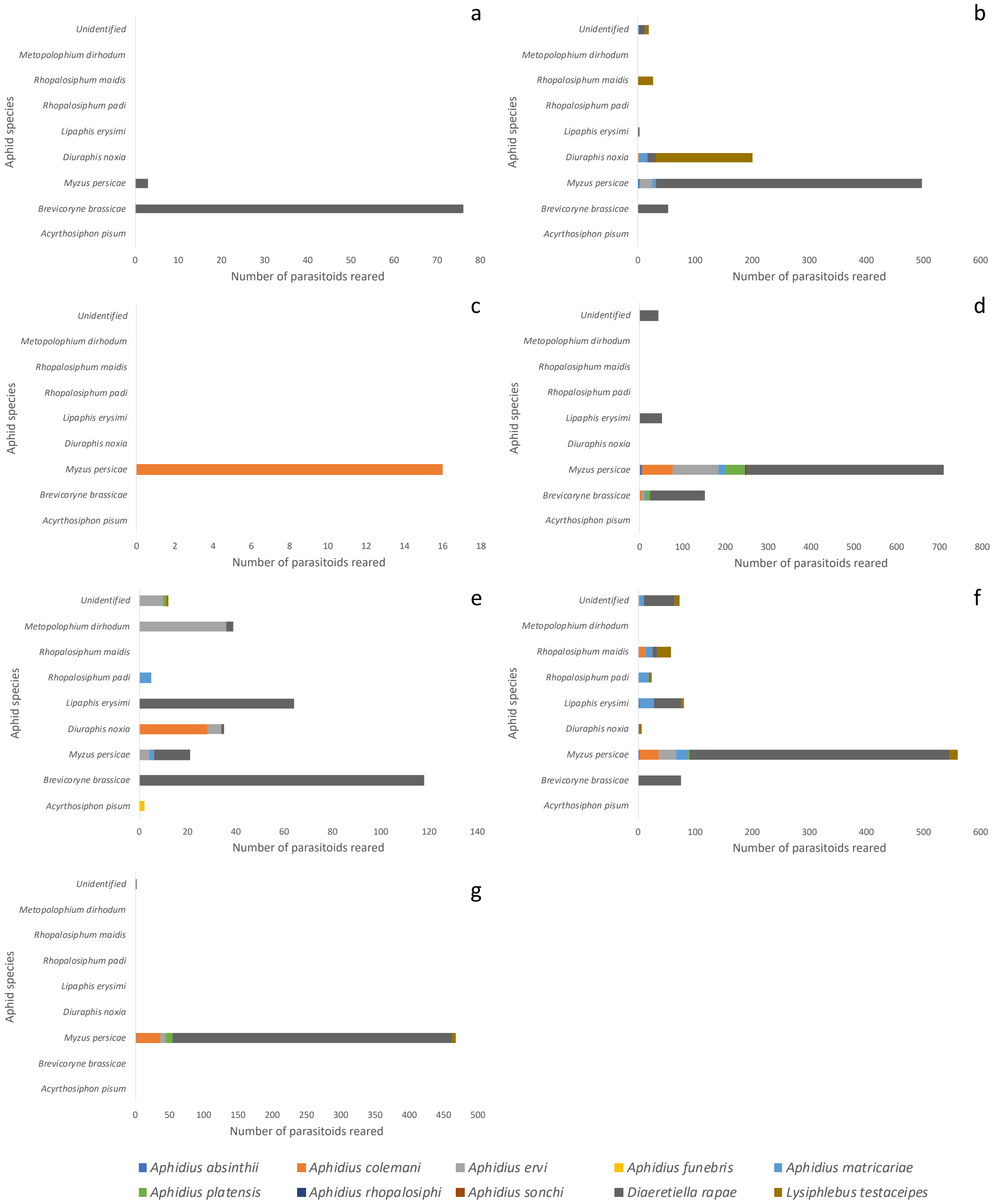


*Figure S4*: Primary parasitoid species composition reared from aphid species in grain production landscapes within ACT (a), NSW (b), QLD (c), SA (d), TAS (e), VIC (f), and WA (g).

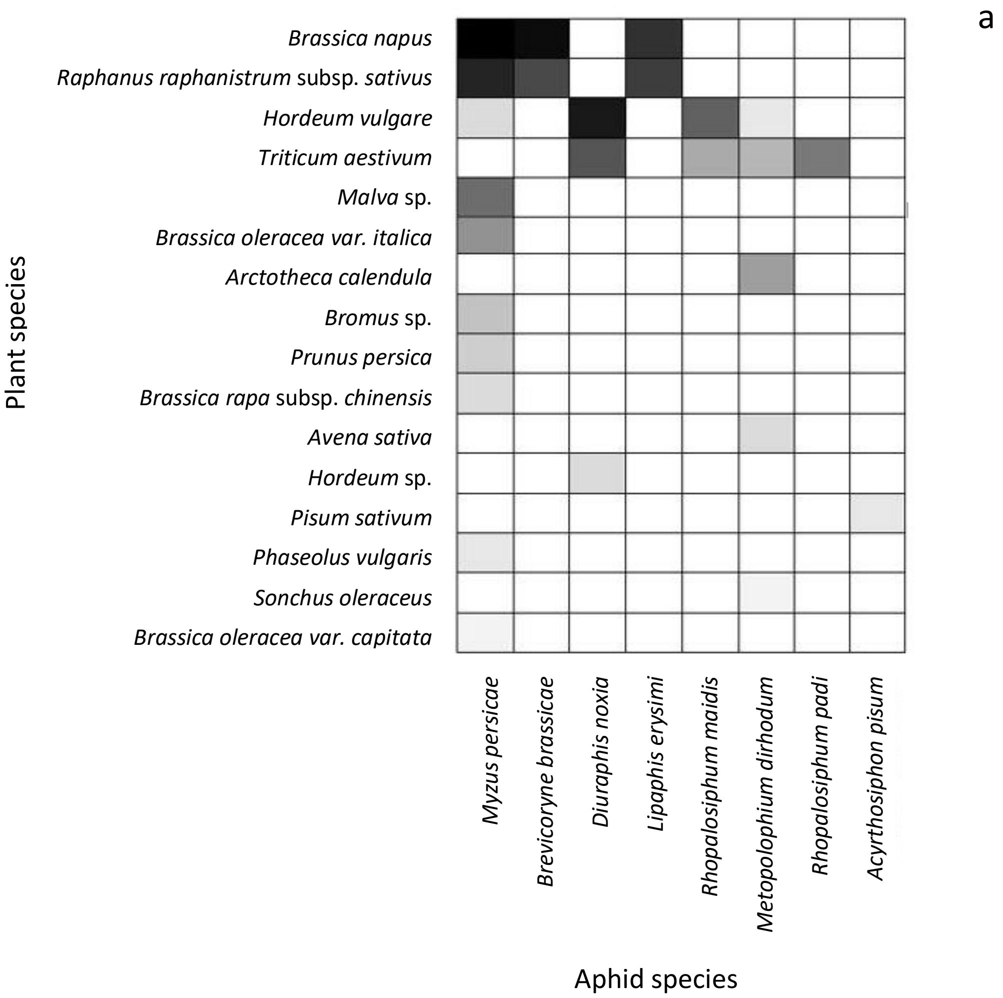


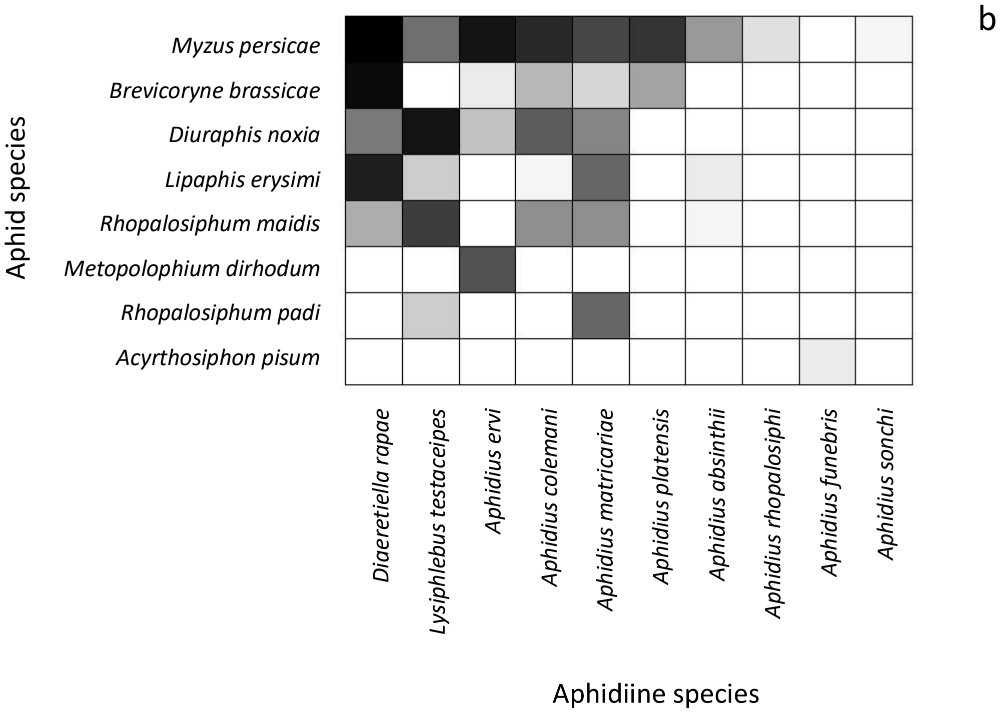


*Figure S5:* Matrices representing the associations observed between a) plants and aphids, and b) aphids and parasitoids, from rearings. [Shade indicates number of interactions, with darker squares indicating more plant-aphid or aphid-wasp interactions occurred, by rearing].
